# The sunlit microoxic niche of the archaeal eukaryotic ancestor comes to light

**DOI:** 10.1101/385732

**Authors:** Paul-Adrian Bulzu, Adrian-Ştefan Andrei, Michaela M. Salcher, Maliheh Mehrshad, Keiichi Inoue, Hideki Kandori, Oded Beja, Rohit Ghai, Horia L. Banciu

**Affiliations:** Department of Molecular Biology and Biotechnology, Faculty of Biology and Geology, Babeş-Bolyai University, Cluj-Napoca, Romania; Institute of Hydrobiology, Department of Aquatic Microbial Ecology, Biology Centre of the Academy of Sciences of the Czech Republic, České Budějovice, Czech Republic.; Limnological Station, Institute of Plant and Microbial Biology, University of Zurich, Seestrasse 187, CH-8802 Kilchberg, Switzerland.; Faculty of Biology, Technion Israel Institute of Technology, Haifa, Israel.; The Institute for Solid State Physics, The University of Tokyo, Kashiwa, Japan; Department of Life Science and Applied Chemistry, Nagoya Institute of Technology, Nagoya, Japan.; Molecular Biology Center, Institute for Interdisciplinary Research in Bio-Nano-Sciences, Babeş-Bolyai University, Cluj-Napoca, Romania

**Author notes:** These authors contributed equally to this work (Paul-Adrian Bulzu, Adrian-Ştefan Andrei). Corresponding author: Rohit Ghai, Institute of Hydrobiology, Department of Aquatic Microbial Ecology, Biology Centre of the Academy of Sciences of the Czech Republic, Na Sádkách 7, 370 05, České Budějovice, Czech Republic.

## Abstract

Recent advances in phylogenomic analyses and increased genomic sampling of uncultured prokaryotic lineages brought compelling evidence in support of the emergence of eukaryotes from within the Archaea domain of life. The discovery of Asgardaeota archaea and their recognition as the closest extant relative of eukaryotes fuelled the revival of a decades-old debate regarding the topology of the tree of life. While it is apparent that Asgardaeota encode a plethora of eukaryotic-specific proteins (the highest number identified to date in prokaryotes), the lack of genomic information and metabolic characterization has precluded inferences about their lifestyles and the metabolic landscape that may have favoured the emergence of the hallmark eukaryotic subcellular architecture. Here, we use advanced phylogenetic analyses to infer the deep ancestry of eukaryotes and genome-scale metabolic reconstructions to shed light on the metabolic milieu of the closest archaeal eukaryotic ancestors discovered till date. In doing so, we: i) generate the largest Asgardaeota genomic dataset available so far, ii) describe a new clade of rhodopsins encoded within the recovered genomes, iii) provide unprecedented evidence for mixotrophy within Asgardaeota, iv) present first-ever proofs that the closest extant archaeal relatives to all eukaryotes (Heimdallarchaeia) have microoxic lifestyles with aerobic metabolic pathways unique among Archaea (i.e. kynurenine pathway) and v) generate the first images of Asgardaeota.

## Main

At the dawn of genomics, the eukaryotes were recognized as amalgamated genetic jigsaws that bore components of both bacterial and archaeal descent^1,2^. This genomic chimerism served as a source of speculation and debate over the nature of the protoeukaryote ancestors^3–5^ and inspirited a plethora of hypothetical scenarios for the processes that led to eukaryogenesis^2,6^. Even though, in light of recent research, eukaryotes came into existence through the interplay between an archaeal host^7^ and a bacterial endosymbiont^8^, the metabolic milieu of the ancestral prokaryotic lineages and their phylogenetic blueprint still remain elusive. Here, we bridge state-of-the-art cultivation-independent genomics, sensitive molecular phylogenetic analyses and genome-scale metabolic reconstructions in order to shed light upon the deep archaeal ancestors of eukaryotes, as well as the metabolic landscape that favored the rise of the ‘nucleated cellular architecture’. The genomic catalogue generated during this study enabled us to confidently resolve the backbone of the Asgardaeota superphylum^9^ (the closest archaeal relatives of eukaryotes described to date) and to narrow down the eukaryotes’ branching point within the tree of life. Collectively, our analyses revealed that contradictory to current opinions^10^ the archaeal protoeukaryote ancestor was likely microaerophilic and possessed the ability to harness the Sun’s energy *via* rhodopsins, and whose mixotrophic metabolism had already acquired unprecedented (within archaea) circuitries for *de novo* aerobic NAD^+^ synthesis.

### Asgardaeota phylogenomics

Homology-based searches were employed to recover Asgardaeota-related contigs from *de novo* metagenomic assemblies of two deep-sequenced lake sediment samples (with contrasting salinities). By utilizing a hybrid binning strategy and performing manual inspection and data curation, we obtained thirty-five Asgardaeota MAGs (metagenome-assembled genomes), spanning three (out of 4) evolutionary lineages within the superphylum: Lokiarchaeia (23), Thorarchaeia (10), and Heimdallarchaeia (2). To the best of our knowledge, by accurately binning 6 026 contigs (total length 55.75 Mbp, average contig length 9 252.5 bp) we generated the largest genomic dataset available to date for this superphylum (in contrast all publicly available MAGs amount to 47.2 Mbp). Due to the challenges associated with reconstructing the evolutionary relationships between archaea and eukayotes^9^, in our inferences we used only those MAGs (n = 8; **Supplementary Table S1**) that harbored at least 75% of total phylogenetic markers (See **Supplementary Table S3**). The maximum-likelihood phylogenetic tree, based on concatenation of small (SSU) and large (LSU) ribosomal RNA genes, pictured for the first time a topology in which eukaryotes branched with high-support from within Asgardaeota (archaea) (Supplementary Figure S1a). Even more remarkably, in addition to recreating a previously described Asgardaeota/Eukaryota branching pattern^9^, we provide unprecedented support for a close evolutionary linkage between Heimdallarchaeia (Asgardaeota superphylum) and eukaryotes (SH-aLRT=97.5; UFBoot=100) (Supplementary Figure S1a). The genome-focused phylogeny of Asgardaeota revealed a pattern of ancestry, divergence, and descent, in which Heimdallarchaeia comprises the basal branch of the superphylum and Thor-/Odinarchaeia the youngest one (Figure 1a). Although dissimilar in branching pattern with the SSU + LSU tree, the phylogenomic one was found to be robust (Figure 1a) and to support a topology brought into attention by an earlier study^9^. The SR4-recoded^11^ Bayesian tree (maxdiff=0.1) resolved with high support (PP=1) the monophyly of Asgardaeota/Eukaryota, but failed to confidently resolve the internal topology of the superphylum and the branching point of eukaryotes (Figure 1b). Noteworthy, in both SR4-recoded Bayesian (Figure 1b) and maximum-likelihood phylogenies (Supplementary Figure S1b) the eukaryotes caused branch instability for Heimdallarchaeia, which was attracted without statistical support within the superphylum (PP=0.52; SH-aLRT=92.4 and UFBoot=88). To further substantiate the phylogenetic connection between Asgardaeota members and eukaryotes, we screened all the recovered MAGs and the publicly available ones (14) for the presence of potential eukaryotic signature proteins (ESP). Similar to previous reports^3,9,12^, the MAGs were found to be highly enriched with ESP (Figure 1c), which further reinforced their ancestral linkage to eukaryotes. In addition to the reported ESP^9^, we identified a potential subunit of the COPII vesicle coat complex (associated with intracellular vesicle traffic and secretion) in Thorarchaeia and proteins that harbor the N-terminal domain of folliculin - a eukaryote-specific protein which is known to be involved in membrane trafficking in humans^13^ (Figure 1c) in Lokiarchaea. Furthermore, we retrieved conclusive hits for the ESP-related domains Ezrin/radixin/moesin C-terminal domain and active zone protein ELKS in Lokiarchaeia.

**Figure 1.**
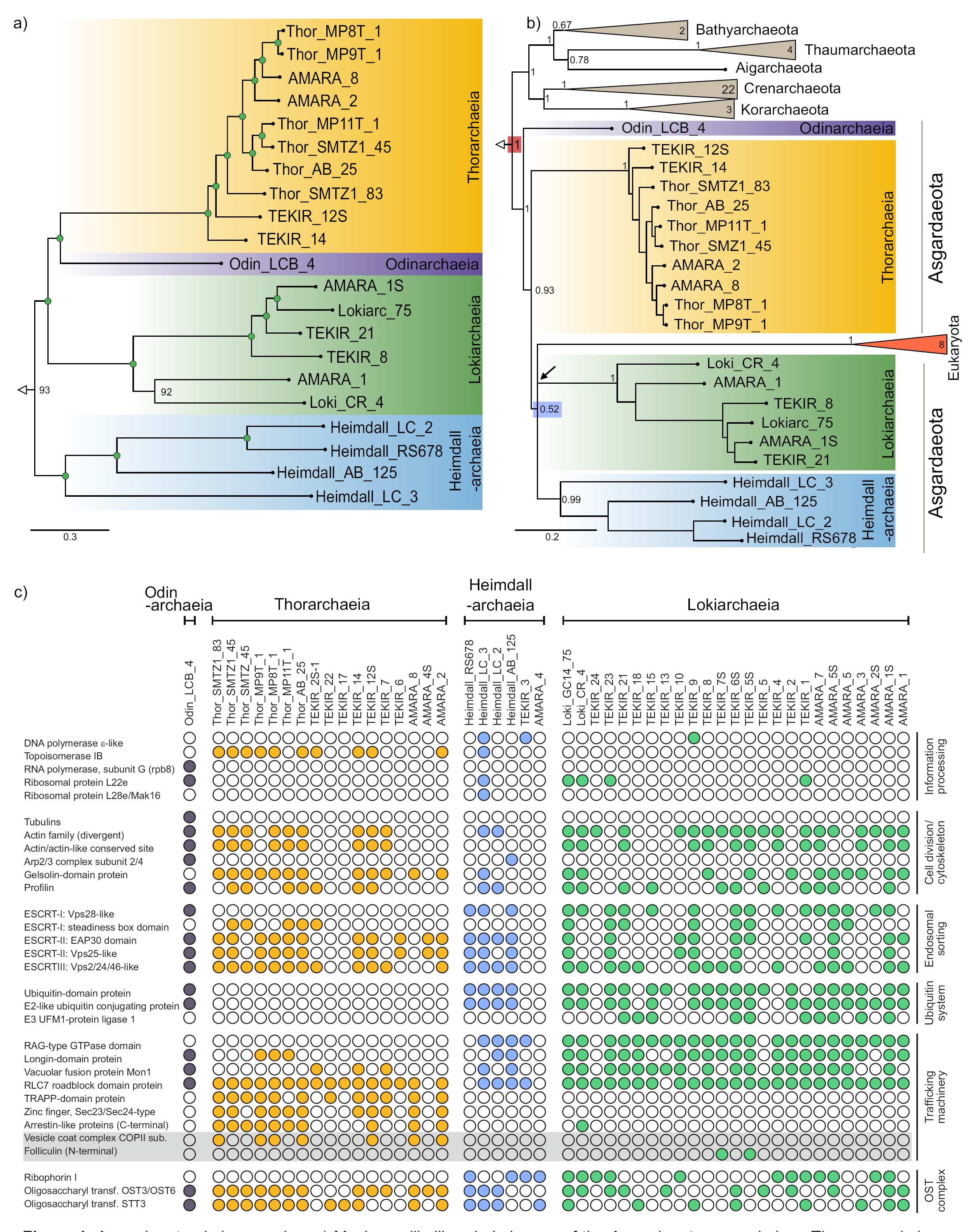
Asgardaeota phylogenomics. a) Maximum-likelihood phylogeny of the Asgardaeota superphylum. The green circles highlight UFBoot values higher than 95. b) Asgardaeota phylogeny generated through Bayesian inference. The posterior probability values are shown above the internal nodes. High support for Eukaryota/Asgardaeota monophyly, and the low support for Eukaryota/Heimdallarchaeia association is indicated by red and blue rectangles on the nodes respectively. The black arrow indicates the unresolved position of Lokiarchaeia. Scale bars indicate the number of substitutions per site. Panel (c) provides a census of the eukaryotic signature proteins (ESP) found in all available MAGs. The grey box highlights new ESPs identified during this study.

### A novel clade of rhodopsins

Recent findings reporting the presence of a novel family of rhodopsins^14^ (i.e. heliorhodopsins; abbreviated as HeR) in monoderms^15^ encouraged us to perform a dedicated screening in all available Asgardaeota MAGs. The results indicated that one of the Heimdallarchaeia MAGs (i.e Heimdallarchaeota RS678) encoded two heliorhodopsins and what appears to be, as suggested by the presence of a *Exiguobacterium*-like DTK motif^16^ and phylogenetic proximity, a type-1 proton-pumping rhodopsin (Figure 2). To the best of our knowledge, this is the first report of a proton-pumping rhodopsin in Asgardaeota. Remarkably, we found that the Asgardaeota MAGs recovered during this study encoded rhodopsin sequences similar in membrane orientation to type-1 rhodopsins, and which organized during phylogenetic analysis in a monophyletic clade (SBS=1) placed in-between HeR and type-1 ones (Figure 2). Multiple sequence alignments showed: i) homology between transmembrane helices 1, 6 and 7 of these new rhodopsins and the type-1 rhodopsins, while helix 3 was homologous to HeR and ii) presence of additional characteristic HeR motifs (e.g. RWxF motif similar to RWxE of HeR rather than the RYxD motif in most type-1 rhodopsins, and replacement of a proline residue (P91) conserved in type-1 rhodopsins by serine (S91) in both HeR and the new Asgardaeota-found ones) (Supplementary Figure S2). Given their phylogenetically intermediate position, as type-1 rhodopsins closest to HeR, and presence of features found in both type-1 and HeR, we denote them as schizorhodopsins (schizo, ‘split’, plus ‘rhodopsin’, abbreviated as SzR). The very recent discovery of HeR and their inconclusive functional role^14,15^ precludes tentative functional assertions for SzR capacity in Asgardaeota. However, the plethora of rhodopsins that we identified in Heimdallarchaeia (putative type-1 proton pumps, HeR and SzR), together with the SzR found in Lokiarchaeia and Thorarchaeia suggests that, during its evolutionary history, Asgardaeota was present in light-exposed habitats. Moreover, we consider that the primary niche of these rhodopsin-bearing microbes is most likely the top, light exposed sediment layers. Their recovery from deeper strata may be explained by the high deposition rates characteristic for the sampling locations (typically a few cm per year)^17^.

**Figure 2.**
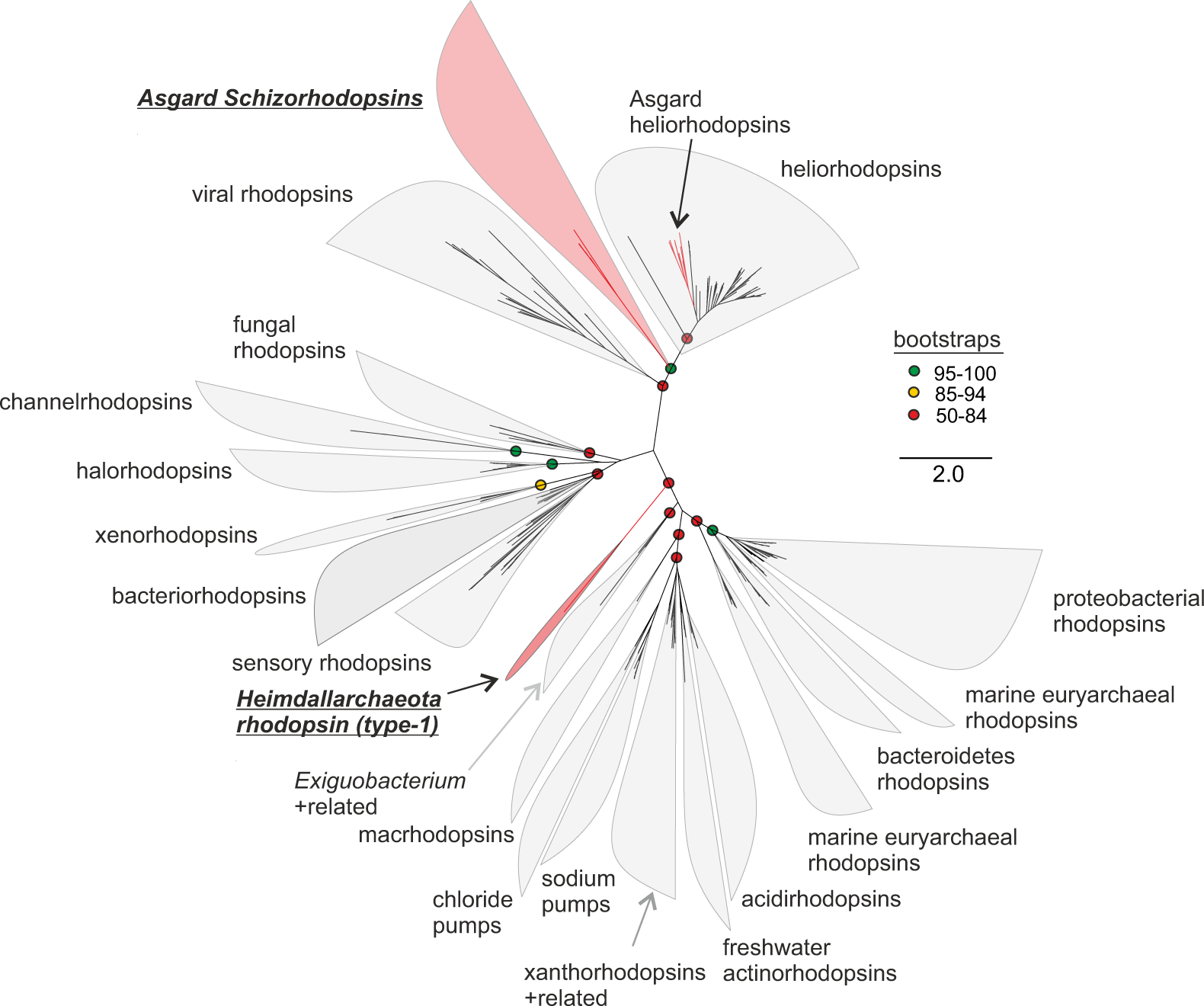
Phylogenetic analysis of rhodopsins. An unrooted maximum-likelihood tree of all Asgardaeota schizorhodopsins identified in this work, heliorhodopsins and representative known type-1 rhodopsins, is shown. The branches colored red are sequences from the Asgardaeota. Bootstrap values on nodes are indicated by colored circles (see color key at the right).

### Evidences for an aerobic lifestyle in Heimdallarchaeia

The genome-scale metabolic reconstruction placed special emphasis on Heimdallarchaeia, since it was suggested by the above-mentioned phylogenetic analyses to encompass the most probable candidates (to date) for the archaeal protoeukaryote. While the anaerobic lifestyles inferred for Loki-^10^ and Thorarchaeia^12^ were considered to be accompanied by autotrophy^10^ and respectively mixotrophy^12^, no consistent metabolic reconstructions exist to date for Heimdallarchaeia. The performed physiological inferences pointed towards mixotrophic lifestyles (for Asgardaeota), simultaneously showing the presence of transporters for the uptake of exogenous organic matter and the metabolic circuitry responsible for its catabolism (see Supplemental Results and Discussion). Remarkably, we found oxygen-dependent metabolic pathways in Heimdallarchaeia, which will be further presented in contrast to the ones harbored by the anaerobic Loki- and Thorarchaeia.

Heimdallarchaeia were inferred to possess components of the aerobic respiration blueprint: a complete tricarboxylic acid cycle (TCA) supported by an electron transport chain (ETC) containing: V/A-type ATPase, succinate dehydrogenase, NADH:quinone oxidoreductase, and the cytochrome c oxidase (Figure 3). While in Thor-various components of the TCA were found to be missing, in Lokiarchaeia the complete TCA was associated with: isocitrate dehydrogenases, 2-oxoglutarate-ferredoxin oxidoreductases, and ATP-citrate lyases, pointing towards the presence of a reverse tricarboxylic acid cycle (rTCA). Thus, in contrast to Heimdallarchaeia, which utilize TCA to fuel their catabolic machinery (Figure 3), Lokiarchaeia uses rTCA for autotrophic CO_2_ assimilation. While the V/A-type ATPase complex appears to be complete in Loki- and Thorarchaeia, the other components involved in the oxidative phosphorylation processes were not identified.

**Figure 3.**
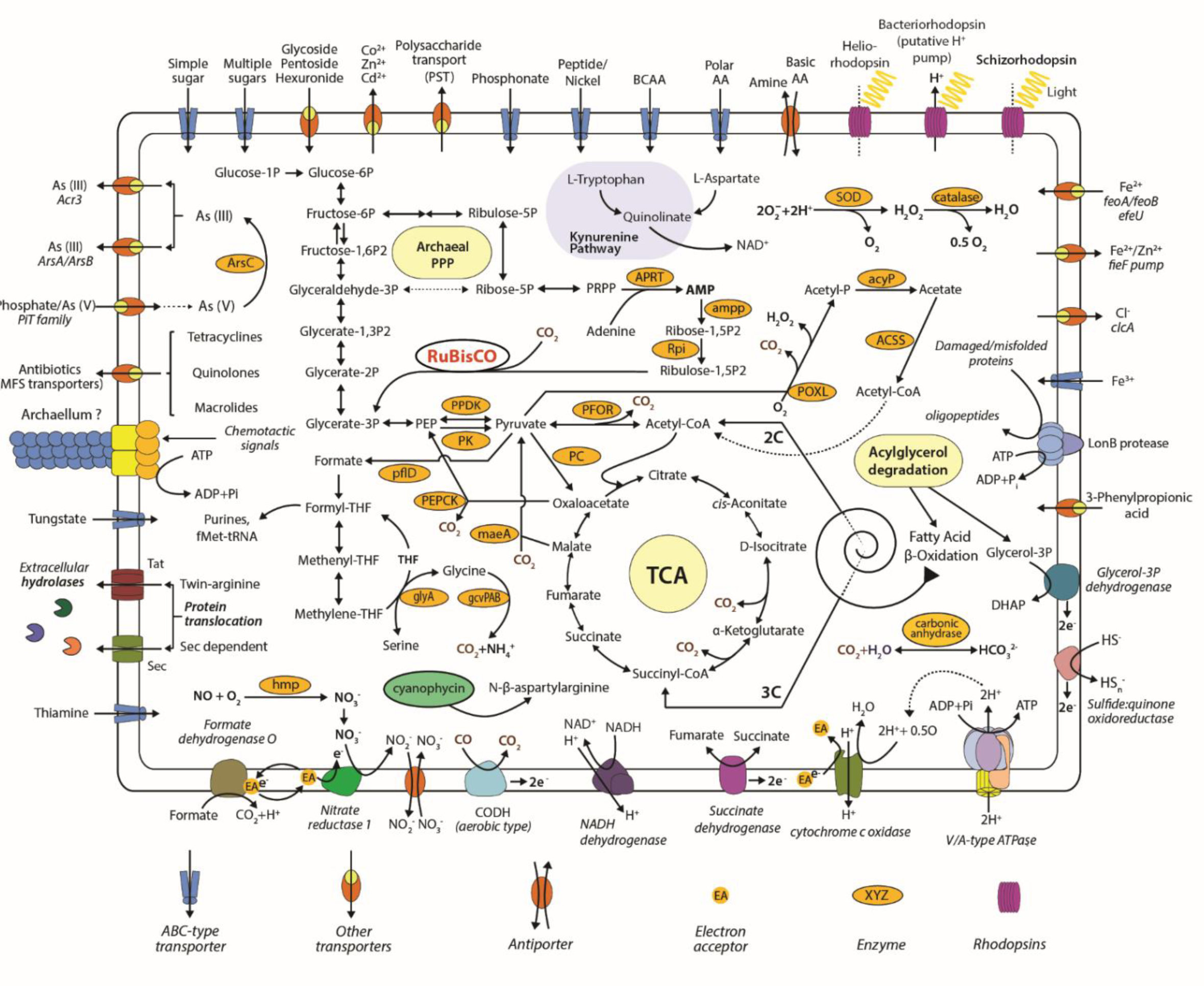
Metabolic reconstruction of Heimdallarchaeia. The text present in the yellow panels depicts names of pathways and metabolic processes. Abbreviations: ArsC - arsenate reductase (glutaredoxin); hmp - nitric oxide dioxygenase; PPDK - pyruvate, phosphate dikinase; PK - pyruvate kinase; PEPCK - phosphoenolpyruvate carboxykinase; maeA - malate dehydrogenase (decarboxylating); PC - pyruvate carboxylase; glyA - glycine hydroxymethyltransferase; gcvPAB - glycine dehydrogenase; hmp - nitric oxide dioxygenase; PFOR - pyruvate ferredoxin oxidoreductase; APRT - AMP pyrophosphorylase; ampp - AMP phosphorylase; Rpi - ribose-5-phosphate isomerase; SOD – superoxide dismutase; catalase; acyP - acylphosphatase; POXL - pyruvate oxidase; ACSS - acetyl-CoA synthetase and carbonic anhydrase. RuBisCO - Ribulose-1,5-bisphosphate carboxylase/oxygenase is shown in red.

As nicotinamide adenine dinucleotide (NAD^+^) is an essential cofactor in redox biochemistry and energetics^18^ (e.g. linking TCA and ETC), we investigated its *de novo* synthesis mechanisms (Figure 4). As expected, all Asgardaeota phyla were found to harbor the aspartate pathway^19^- a set of metabolic transformations that can occur in both presence or absence of oxygen^20^, and which are characteristic for most prokaryotes and the plastid-bearing eukaryotes (obtained through lateral gene transfer from their cyanobacterial endosymbiont)^18^. Surprisingly, Heimdallarchaeia presented in addition to the aspartate pathway the exclusively aerobic kynurenine one^21^ (Figure 4), which is reported to be present in few bacterial groups and eukaryotes^18^. The phylogenetic reconstruction and evolutionary history inferences showed that this pathway, which is considered to be present in the protoeukaryote ancestor^18^, was probably acquired by Heimdallarchaeia through lateral gene transfer from bacteria (Supplementary Figure S3). As far as the authors are aware, Heimdallarchaeia are the first archaeal organisms with the aerobic kynurenine pathway. Curiously while Heimdall_LC_3 was found to contain the complete set of genes required for both pathways, Heimdall_LC_2 and Heimdall_RS678 encoded just the genes affiliated with the kynurenine one (Figure 4). As the aspartate pathway was reported to function in both oxygen absence (L-aspartate oxidase uses fumarate as electron acceptor)^20^ and presence (L-aspartate oxidase uses O_2_ as electron acceptor), the existence of the kynurenine pathway in Heimdall_LC_3 appears redundant. By corroborating the presence/absence pattern of the aspartate pathway in Asgardaeota (Figure 4) with the reconstructed evolutionary history of Heimdallarchaeia (Figure 1a, b; Supplementary Figure S1a, b) and blastp similarity searches (for Heimdall_LC_3 L-aspartate oxidase), we inferred that this pathway functioned exclusively under anaerobic conditions. Furthermore, the introgression of kynurenine genes in Heimdallarchaeia appears to be caused by an expansion towards an oxygen-containing niche, which during evolutionary history (from Heimdall_LC_3 to Heimdall_LC_2/Heimdall_RS678) favored the xenologous replacement of the aspartate pathway with the kynurenine one.

**Figure 4.**
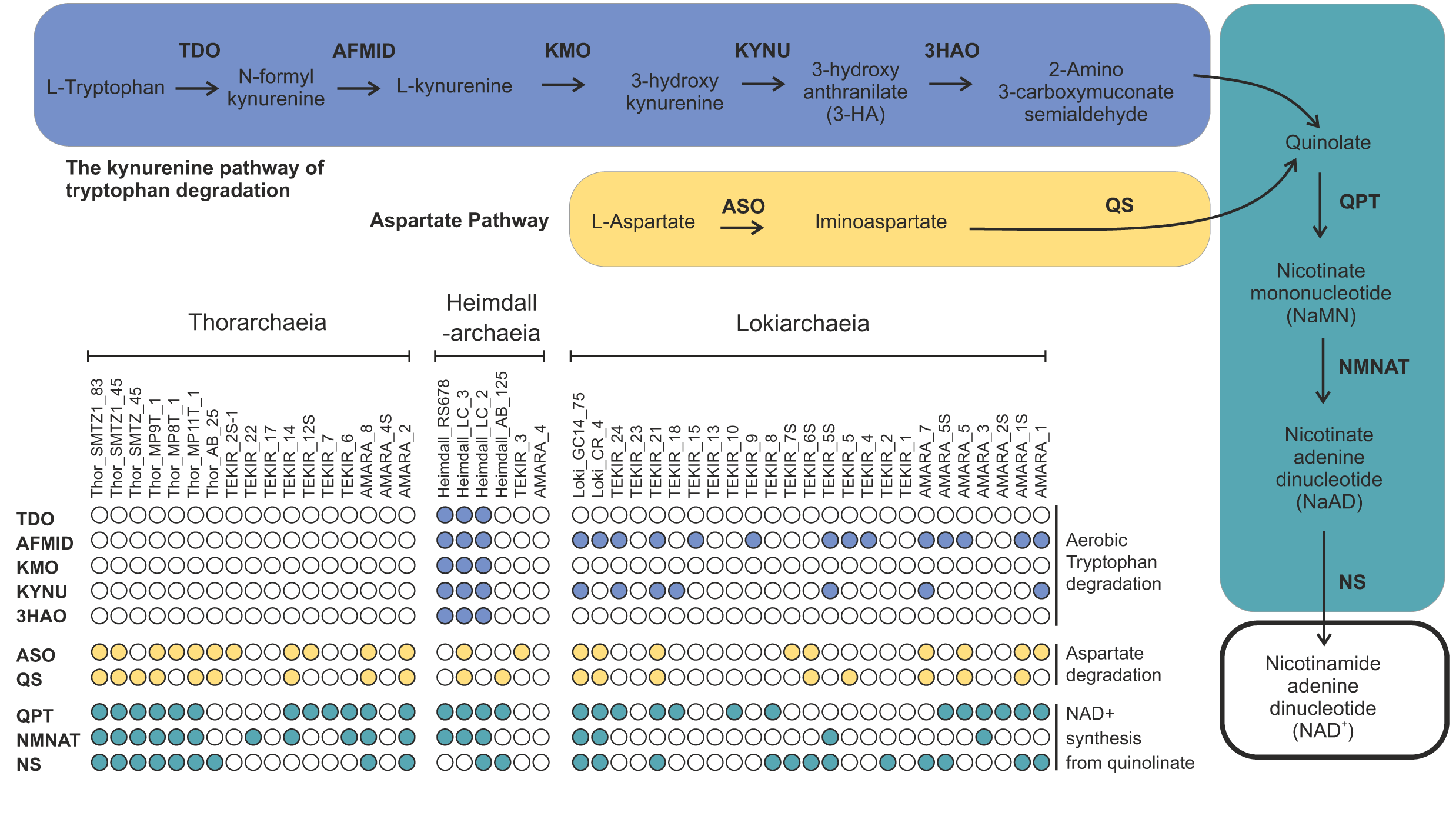
*De novo* NAD^+^ synthesis pathways. The colored boxes show a schematic representation of the kynurenine and aspartate pathways involved in *de novo* NAD^+^ synthesis. The presence of the enzymes involved in these pathways is indicated for each MAG by using a colored circle. TDO - tryptophan 2,3-dioxygenase; AFMID – arylformamidase; KMO - kynurenine 3- monooxygenase; KYNU – kynureninase; 3HAO - 3-hydroxyanthranilate 3,4-dioxygenase; ASO - L-aspartate oxidase; QS - quinolinate synthase; QPT - nicotinate-nucleotide pyrophosphorylase; NMNAT - nicotinamide-nucleotide adenylyltransferase; NS - NAD^+^ synthase

Within the anaplerotic metabolism, the reversible transformation of pyruvate into acetyl-CoA and formate can be accomplished by pyruvate formate lyases, which were inferred to be present in all three phyla. Formate produced during this enzymatic process, or by the activity of arylformamidase (kynurenime formamidase) in Loki- and Heimdallarchaeia could be further oxidized (by formate dehydrogenases) and used for quinone/cytochrome pool reduction, or introduced into the one-carbon metabolism and utilized for the synthesis of: purines, glycine, formylmethionine, etc. (**Figure** 3). Uniquely in Heimdallarchaeia we inferred that formate could act as electron donor during aerobic respiration through the actions of the heterotrimeric formate dehydrogenase O. This enzyme facilitates the usage of formate under aerobiosis, and together with nitrate reductase Z (also present solely in Heimdallarchaeia) may participate in a formate to nitrate electron transport pathway that is active when cells are shifted from aerobic to anaerobic conditions^22,23^. The presence of genes encoding pyruvate oxidases (poxL) in Heimdallarchaeia (i.e. LC_2 and LC_3) implies further oxygen usage, as the enzyme employs it in the pyruvate pool decarboxylation process (Figure 3).

The comparative genomic analyses also revealed that the three Asgardaeota phyla rely upon glycolysis (i.e. type Embden-Meyerhof-Parnas) to fuel their metabolic machinery. Unexpectedly, three Heimdallarchaeia MAGs (LC_3, AB_125 and AMARA_4) were found to employ non-canonical ADP-dependent kinases that use ADP instead of the typical ATP as the phosphoryl group donor^24^ in their glycolytic pathways. Furthermore, they seemed to be bifunctional ADP-dependent glucokinase/phosphofructokinases, which was puzzling since the presence of 6- phosphofructokinases (LC_3 and AB_125) would render their bifunctionality redundant. In order to elucidate the role of the putative bifunctional enzymes, we reconstructed the evolutionary history of the ADP-dependent kinases and inferred that they possess glucokinase activity (based on tree topology and the conserved functional residue E172) (Supplementary Figure S4). Additionally, we observed that the deepest branching Heimdallarchaeia (LC_3) harbored the archaeal-type enzyme, while the younger ones (AB_125 and AMARA_4) clustered together with the eukaryotic-type (Supplementary Figure S4). Although it is easy to assume that cells under low energy conditions (e.g. limiting O_2_ availability) could highly benefit from using residual ADP to activate sugar moieties and fuel their glycolysis^25^, the metabolic advantage conferred by these ADP-dependent kinases is unclear.

Although pentoses could be recycled via nucleotide degradation in all Asgardaeota phyla, their synthesis differs between Loki-/Heimdallarchaeia that likely utilize the reverse ribulose monophosphate pathway, and Thorarchaeia that employ the xylulose part (of the non-oxidative branch) of the hexose monophosphate one. The identified homologues for ribulose 1,5-bisphosphate carboxylase/oxygenase (RuBisCO) genes were found to appertain to the types: III (Loki- and Heimdallarchaeia) and IV (Loki- and Thorarchaeia) (Supplementary Figure S5). While RuBisCO is a key enzyme for CO_2_ fixation in the Calvin-Benson-Bassham cycle, the absence of phosphoribulokinase renders this metabolic pathway highly improbable. However, we consider that the MAGs encoding type III-like RuBisCO (assigned to Loki- and Heimdallarchaeia) use the nucleotide monophosphate degradation pathway^26,27^, thus performing CO_2_ fixation by linking nucleoside catabolism to glycolysis/gluconeogenesis. This conclusion is supported by the co-occurrence of genes encoding for: RuBisCO type III, AMP phosphorylases, ribose 1,5-bisphosphate isomerases, and carbonic anhydrases. While carbon monoxide (CO) can be used as carbon and energy source in both aerobic and anaerobic metabolisms^28^, the types of enzymes involved in the reaction are dependent upon the available electron acceptor. Thus, while Heimdallarchaeia harbor all three major subunits of the aerobic carbon monoxide dehydrogenases (CODH), Loki- and Thorarchaeia encoded the oxygen-sensitive carbon monoxide dehydrogenase/acetyl-CoA synthase (CODH/ACS). We infer that while Heimdallarchaeia uses CO to obtain energy by shuttling the electrons generated from CO oxidation to oxygen or nitrate, Thor- and Lokiarchaeia may utilize CO as both electron source and intermediary substrate in the ancient Wood–Ljungdahl carbon fixation pathway^29,30^ (through CODH/ACS).

Among Asgardaeota, the Heimdallarchaeia were found to possess genes encoding for sulfide:quinone oxidoreductases, enzymes used in sulfide detoxification and energy generation via quinone pool reduction (Figure 3). As sulfide binds to cytochrome c oxidase system and inhibits aerobic respiration^31^, the presence of these enzymes in Heimdallarchaeia could point towards a detoxification role. The fact that one Heimdallarchaeia MAG described in this study (i.e. AMARA_4) had the sulfide:quinone oxidoreductase gene and the other Asgardaeota MAGs recovered from the same sample did not (i.e. 7 Loki- and 3 Thorarchaeia MAGs), suggests that the highly lipophilic sulfide does not interfere with the anaerobic metabolism, nor is it part of a conserved energy generation strategy in Asgardaeota. The superoxide dismutase, catalases, and glutathione peroxidases found in Heimdallarchaeia may act in alleviating the oxidative damaged generated by a facultative aerobic metabolism.

### CARD-FISH visualization of Loki- and Heimdallarchaeia

Two phylogenetic probes targeting the 16S rRNA of Loki- and Heimdallarchaeia, respectively (**Supplementary Table S8, Supplementary Figure S7**) were successfully applied to sediment samples of different depth layers. Members of both phyla were rare and seemed to be totally absent below sediment depths of 40 cm. All observed Heimdallarchaeia were similar in cell size (2.0±0.4 µm length x 1.4±0.3 µm width, n=15) and of conspicuous shape with DNA condensed (0.8±0.2 x 0.5±0.1 µm) at the center of the cells (Figure 5 a-c, **Supplementary Figure S8**), which is rather atypical for prokaryotes. In contrast, Lokiarchaeia were very diverse in shape and size and we could distinguish at least three different morphotypes: small-medium sized ovoid cells (2.0±0.5 x 1.3±0.3 µm, n=23, Figure 5 d-f, **Supplementary Figure S9**), large round cells (3.8 x 3.6 µm, Figure 5 g-i) with condensed DNA at the center, and large rods/filaments (4.4±1.2 x 1.4±0.5 µm, n=6, Figure 5 j-l, **Supplementary Figure S9**) with filamentous, condensed DNA (2.7±1.4 x 0.4±0.1 µm) that were exclusively present in 30-40 cm sediment depth. This high diversity in morphology in Lokiarchaeia most likely also reflects a higher sampling of the phylogenetic diversity within the phylum.

**Figure 5.**
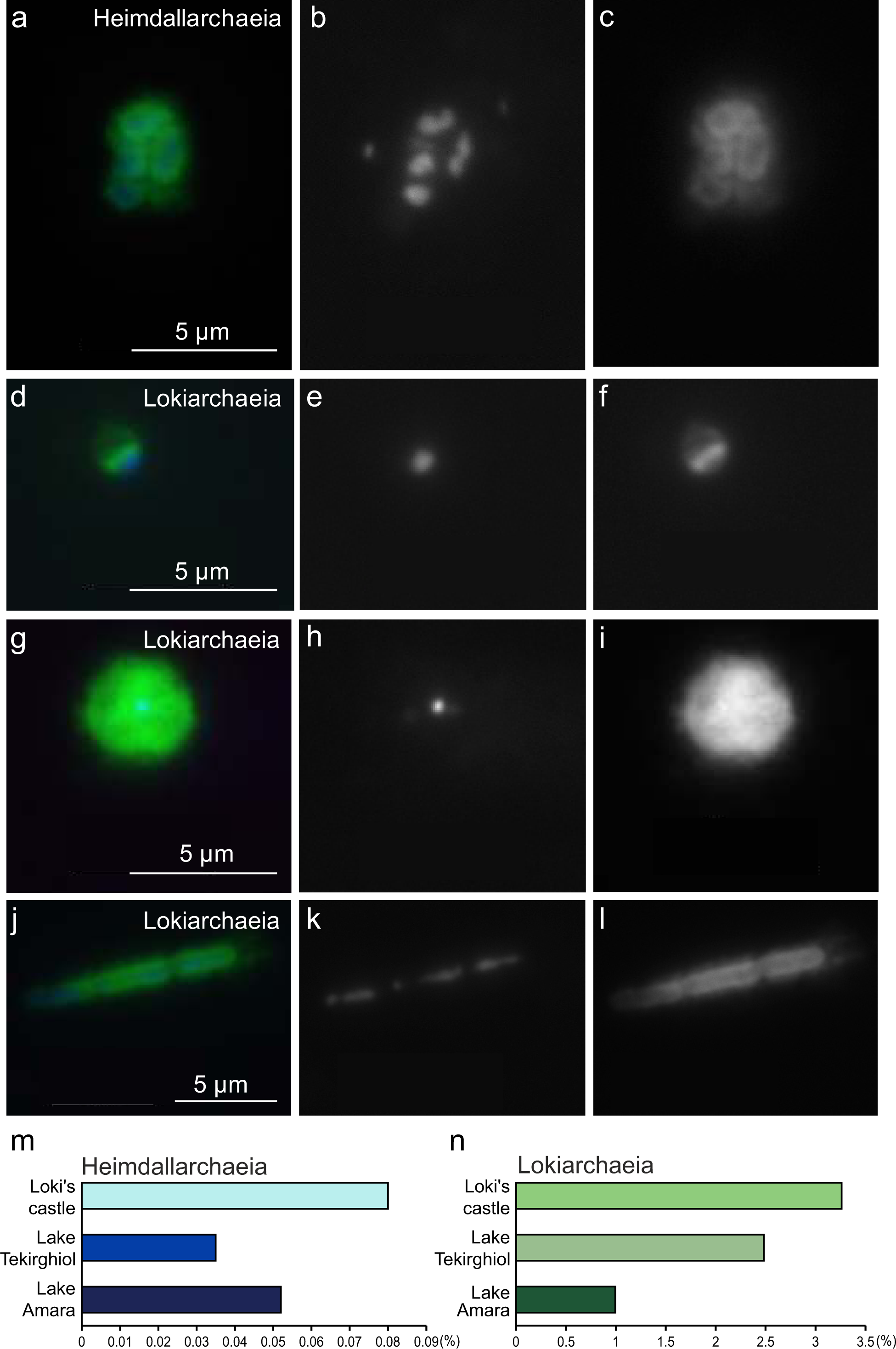
CARD-FISH imaging of Heimdallarchaeia and Lokiarchaeia. The upper panel (a-c) shows several cells of Heimdallarchaeia hybridized with probe heimdall-526, the lower three panels (d-l) diverse morphologies of Lokiarchaeia targeted by probe loki1-1184. The left panels (a, d, g, j) display overlay images of probe signal (green), DAPI staining (blue) and autofiuorescence (red), the middle panels (b, e, h, k) DAPI staining of DNA, the right panels (c, f, i, l) CARD-FISH staining of proteins. Abundance estimation of recovered 16S rRNA reads afiliated with Heimdallarchaeia (m) and Lokiarchaeia (n) within metagenomes from Loki’s castle, Lake Tekirghiol, and Lake Amara.

## Discussion

The mainstream theories on the subject of eukaryogenesis^6,32^ (which date back to late 20^th^ century), have been recently refuted by improved phylogenetic methods and increased genomic sampling^9,10^. Even after experiencing a new revival^10^, the current endosymbiotic theory fails to envision the environmental and metabolic context in which the ‘nucleated cellular architecture’ emerged. Moreover, it appears to find itself gravitating around a three-decade old theory centered on anaerobic syntrophy^33^ (i.e. hydrogen hypothesis). In this study, we show that: i) mixotrophy and harnessing the Sun’s energy (*via* rhodopsins) are the *modus vivendi* of Asgardaeota, and that ii) aerobic respiration was present in the archaeal protoeukaryote ancestor before the emergence of eukaryogenesis-associated phenomena. The ‘aerobic-protoeukaryotes’ model surpasses some of the theoretical shortcomings of the ‘hydrogen hypothesis’ by envisioning an endosymbiotic association in which the primordial function of the bacterial counterpart (i.e. oxidative phosphorylation) would not be detrimental to the existence of the archaeal host (caused by oxygen exposure). Furthermore, this model is supported by a recent high-scale time-calibrated phylogenomic tree which shows that the archaeal-bacterial endosymbiosis that gave birth to the protoeukaryote ancestor took place after the Great Oxidation Event^34^. In light of these results, we consider that previous hypotheses regarding the metabolic capacities of the protoeukaryote ancestors need to be reevaluated.

## Methods

### Sampling

Sediment sampling was performed on 10 October 2017 at 12:00 in Tekirghiol Lake, Romania, (44°03.19017 N, 28°36.19083 E) and on 11 October 2017 at 15:00 in Amara Lake, Romania, (44°36.30650 N, 27°19.52950 E). Two plunger cores of 0.3 m each were collected from a water depth of 0.8 m in Tekirghiol Lake and 4 m in Amara Lake. Sediment samples were stored in the dark at 4 °C and processed within 24 hours after collection.

### DNA extraction and purification

DNA was extracted from approximately 10 g of wet sediment from each mixed core sample using the DNeasy PowerMax Soil Kit (Qiagen, Hilden, Germany) following the manufacturer’s instructions. Extracted DNA was further purified by passing it through humic acid removal columns (type IV-HRC) provided in the ZR Soil Microbe DNA MiniPrep kit (Zymo Research, Irvine, CA, USA). Purified DNA was quality checked and quantified using a ND-1000 NanoDrop spectrophotometer (Thermo Scientific, Waltham, MA, USA). DNA integrity was assessed by agarose gel (1%) electrophoresis and ethidium bromide staining. The samples were denominated as AMARA and TEKIR in accordance with their site of origin. From each sample, 4 µg of pure DNA were vacuum dried in a SpeedVac concentrator (Thermo Scientific, Waltham, MA, USA) and shipped for library construction and NGS sequencing to Macrogen (Seoul, South Korea).

### Sequencing and data preprocessing

Library preparation was performed by the commercial company by using the TruSeq DNA PCR Free Library prep kit (Illumina). Whole-genome shotgun sequencing of the 150 paired-end libraries (350bp insert size) was done using a HiSeq X (Illumina) platform. The amount of total raw sequence data generated for each metagenome was: 64.5Gbp for Amara and 57.6 Gbp for Tekirghiol. Preprocessing of raw Illumina reads was carried out by using a combination of software tools from the BBMap^35^ project (https://github.com/BioInfoTools/BBMap/). Briefly, bbduk.sh was used to remove poor quality sequences (qtrim=rl trimq=18), to identify phiX and p-Fosil2 control reads (k=21 ref=vectorfile ordered cardinality), and to remove Illumina adapters (k=21 ref=adapterfile ordered cardinality).

### Abundance estimation for Loki- and Heimdallarchaeia

Preprocessed Illumina sets from Amara and Tekirghiol lakes, as well as published^3^ set SRX684858 from Loki’s castle marine sediment metagenome, were subsampled to 20 million reads by reformat.sh^36^. Each subset was queried for putative RNA sequences by scanning with UBLAST^37^ against the non-redundant SILVA SSURef_NR99_132 database^38^, that was priorly clustered at 85% sequence identity by UCLUST^37^. Identified putative 16S rRNA sequences (e-value < 1e-5) were screened using SSU-ALIGN^39^. Resulting bona fide 16S rRNA sequences were compared by blastn^40^ (e-value <1e-5) against the curated SILVA SSURef_NR99_132 database. Matches with identity ≥ 80% and alignment length ≥90 bp were considered for downstream analyses. Sequences assigned to Loki- and Heimdallarcheia were used to calculate abundances for these taxa in their originating environments **(Figure 5).**

### Metagenome assembly and binning

*De novo* assembly of preprocessed paired-end Illumina reads was done by Megahit^41^ v.1.1.1 with k-mer list: 39, 49, 69, 89, 109, 129, 149, and with default parameters. Assembled contigs with minimum nucleotide fragment length of 3 kbp were binned by a combination of taxonomy-dependent and -independent methods. Protein coding genes were predicted by MetaProdigal^42^. Taxonomy dependent binning was achieved by first assigning taxonomy labels to the predicted genes by performing screenings with MMseqs2^43^ against the NR database. All contigs with a minimum of 30 % genes assigned to Asgardaeota were used for taxonomy-independent binning. Mean base coverage for each contig was computed with bbwrap.sh (default parameters) by mapping to assembled contigs the preprocessed reads from AMARA and TEKIR datasets. Hybrid binning (based on tetranucleotide frequencies and coverage data) was performed using MetaBAT2^44^ with default parameters. Bin completeness, contamination and strain heterogeneity were estimated using CheckM^45^ with default parameters. Poorly resolved bins (i.e. contamination > 10%, unbinned contigs) were further manually curated by a combination of tetranucleotide frequency PCA graphs and repeated rounds of contamination/completeness assessment by CheckM. Final curated bins with CheckM estimated completeness above 10% and contamination below 10% were denominated as metagenome assembled genomes (MAGs). A total of 35 MAGs were recovered: 23 Lokiarchaeia, 10 Thorarchaeia and 2 Heimdallarchaeia (**Supplementary Table S1**). Unbinned contigs were kept for further analyses (total nucleotide bases/site: 3.46 Mbp Amara and 4.06 Mbp Tekirghiol).

### Genome annotation

Publicly available Asgardaeota genomes were downloaded from the NCBI Genome section (https://www.ncbi.nlm.nih.gov/genome). Coding sequences were predicted *de novo* with Prokka^46^ for all available Asgard MAGs (35 from this study, 14 from NCBI – Accession numbers can be found in **Supplementary Table S2**). BlastKOALA^47^ was used to assign KO identifiers (K numbers) to orthologous genes (**Supplementary Table S3**). Inferences of metabolic pathways and general biological functions were conducted with the online KEGG mapping tools (https://www.genome.jp/kegg/kegg1b.html) using summarized KO numbers assigned to each group. Odinarchaeia was not considered for metabolic reconstruction due to lack of new genomic data. Ribosomal RNA (rRNA)-coding regions (16S, 23S) and transfer RNA (tRNA)-coding regions were predicted with Barrnap (https://github.com/tseemann/barrnap) and tRNAscan-SE^48^, respectively. All predicted proteins were queried against NCBI NR, COGs (cluster of orthologous groups) and arCOGs (archaeal cluster of orhtologous groups, 2014)^49^. A local version of InterProScan^50^ was used with default settings to annotate protein domains. Potential eukaryote specific proteins (ESPs) were identified based on previously published lists of IPR domains^9^ (**Supplementary Table S4**) identified in Asgard archaea. New ESPs were searched based on key words related to eukaryotic specific processes and/or structures. All IPR domains present exclusively in newly recovered Asgardaeota genomes were manually screened by querying accession numbers against the online InterPro database (https://www.ebi.ac.uk/interpro/search/sequence-search), for associations with eukaryotic specific domains. A previously identified^9^ ESP - DNA polymerase epsilon, catalytic subunit (IPR029703) - was identified by querying all MAG proteomes with human sequences (**Supplementary Table S4**). Several candidate ESP sequences were further analyzed using jackhmmer^51^, Phyre2^52^ and Phobius^53^.

### Phylogenetic trees

A total of 131 taxa were considered for concatenated small subunit (SSU) and larger subunit (LSU) ribosomal RNA phylogenetic analyses, consisting of: 97 archaea (37 Euryarchaeota, 24 Crenarchaeota, 2 Bathyarchaeota, 15 Thaumarchaeota, 3 Aigarchaeota, 2 Korarchaeota, 14 Asgardaeota), 21 bacteria and 13 eukaryotes (**Supplementary Table S5**). SSU and LSU sequences were aligned independently by PRANK^54^ (parameters: -DNA +F), trimmed using BMGE^55^ (–m DNAPAM250:4 –g 0.5) and concatenated. Members of the DPANN group of Archaea (Diapherotrites, Parvarchaeota, Aenigmarchaeota, Nanoarchaeota, Nanohaloarchaeota, Woesearchaeota, and Pacearchaeota) were not included due to their known tendency to cause phylogenetic artefacts (detailed previously^9^). Maximum-Likelihood phylogeny for concatenated SSU/LSU gene sequences was inferred using IQ-TREE (-m GTR+I+G4+F) with ultrafast bootstrapping - bb 1000 and Shimodaira-Hasegawa testing –alrt 1000^56,57^.

A total of 93 taxa were considered for concatenated ribosomal protein phylogenomic analyses, consisting of: 85 Archaea (25 Euryarchaeota, 22 Crenarchaeota, 2 Bathyarchaeota, 4 Thaumarchaeota, 1 Aigarchaeota, 3 Korarchaeota, 21 Asgardaeota, 7 DPANN) and 8 eukaryotes (**Supplementary Table S5**). Selection criteria for phylogenomic trees of ribosomal proteins conserved between archaea and eukaryotes has been previously described^9^. Amino-acid sequences for the 55 ribosomal proteins were queried and retrieved based on arCOG annotations. Markers not found in the majority of organisms were discarded, obtaining a final set of 48 markers (**Supplementary Table S6**). Additionally, some proteins that were not identified by arCOG scanning were retrieved from the NCBI Protein section (https://www.ncbi.nlm.nih.gov/protein). Sequences were aligned using PRANK (-protein +F), trimmed with BMGE^55^ (-m BLOSUM30 -t AA -g 0.2 -b 3), concatenated, and subjected to SR4 amino acid recoding^11^. Maximum-likelihood trees were generated by IQ-TREE (-bb 1000, -alrt 1000) with ultrafast bootstraping^56^ and the custom ‘C60SR4’ model described in a previous study^9^. Bayesian inference phylogenies were constructed using PhyloBayes MPI 1.8^58^, using the CAT-Poisson model. Four chains were run in parallel until estimated maxdiff values calculated by bp_comp (-x 5000 10) fell below the recommended 0.3 threshold, indicating convergence between chains.

### Multiple sequence alignment of rhodopsins

The three groups of rhodopsins (type-1, schizorhodopsins and heliorhodopsins), were first aligned independently using T_Coffee^59^ (http://tcoffee.crg.cat/) in accurate mode, that employs protein structure information, wherever available, or sequence comparisons with homologues in databases to improve accuracy. These alignments were aligned to each other using the profile alignment mode in T_Coffee.

### RuBisCO tree reconstruction

The multiple sequence RuBisCO alignment was built upon a core structural alignment of diverse set of sequences, to which additional sequences were added using T_Coffee. A total of 392 sequences were used for the alignment. MUSCLE^60^ was used for aligning the sequences (n=146) of the large subunit of RuBisCO (types I-III) and RuBisCO-like (type IV) (rbcL, K01601) proteins. Sequences not generated in this study were recovered from previous studies^61,62^. For both alignments the maximum likelihood tree was constructed with FastTree2 using a JTT model, a gamma approximation, and 100 bootstrap replicates.

### Phylogenetic inference of Heimdallarchaeia glucokinases and kynurenine pathway proteins

ADP- dependent phosphofructokinase/glucokinase protein sequences were identified by their assigned KO number (K00918) in 3 MAGs (AMARA_4, Heimdall_AB_125, Heimdall_LC_3). Retrieved sequences were used along with 49 other sugar kinases published in a previous study^63^. Protein sequences of components of the kynurenine pathway - tryptophan 2,3-dioxygenase (TDO), kynurenine 3-monooxygenase (KMO) and 3-hydroxyanthranilate 3,4-dioxygenase (HAAO) – that were identified only in Heimdallarchaeia MAGs, were used along with sequences of corresponding enzymes from 12 Eukaryotes and 15 Bacteria that were retrieved from NCBI RefSeq (Accession numbers in **Supplementary Table S7**). MAFFT-L-INS-i^64^ (default parameters) and PRANK^54^ (parameters: -DNA +F) were used for aligning sugar kinase and respectively kynurenine pathway enzyme sequences followed by trimming using BMGE^55^ (-m BLOSUM30 -t AA -g 0.5 -b 3). Single protein maximum likelihood trees were constructed with FastTree2^65^, using an accurate search strategy (-mlacc 2 –spr 4 –slownni), and 100 bootstrap replicates.

### Probe design and CARD-FISH (Catalyzed reporter deposition fluorescence *in situ* hybridization)

All assembled 16S sequences classified as Asgardaeota were aligned with the SINA aligner^66^, manually optimized in ARB^67^ using SILVA database SSURef_NR99_132^38^, and a RAxML tree (Randomized Axelerated Maximum Likelihood tree with GTR-GAMMA model, 100 bootstraps^68^) was constructed (**Supplementary Figure S7**). Probe design for Heimdallarchaeia and Lokiarchaeia based on almost full-length sequences of high quality was done with the probe_design and probe_check tools in ARB. Probes were tested *in silico*^69^ and in the laboratory with different formamide concentrations in the hybridization buffer until stringent conditions were achieved (**Supplementary Table S8**). Sediment sampling was performed using a custom mud corer on 22 April 2018 at 12:00 in Tekirghiol Lake, Romania, (44°03.19017 N, 28°36.19083 E) and on 23 April 2018 at 14:00, in Amara Lake, Romania, (44°36.30650 N, 27°19.52950 E). Seven sediment layers (0-70 cm, in 10-cm ranges) were sampled in Tekirghiol Lake and the top 10 cm was sampled in Amara Lake. Sediment samples were fixed with formaldehyde, treated by sonication, vortexing and centrifugation to detach cells from sediment particles^70^ and aliquots were filtered onto white polycarbonate filters (0.2 µm pore size, Millipore). CARD-FISH was conducted as previously described with fluorescein labelled tyramides^71^. Filters were counterstained with DAPI and inspected by epifluorescence microscopy (Zeiss Imager.M1). Micrographs of CARD-FISH stained cells were recorded with a highly sensitive charge-coupled device (CCD) camera (Vosskühler) and cell sizes were estimated with the software LUCIA (Laboratory Imaging Prague, Czech Republic).

### Accession numbers

All sequence data produced during the study is deposited in the Sequence Read Archive (SRA) database of the National Center for Biotechnology Information (NCBI) and can be found linked to the Bioproject PRJNA483005.

**End notes.**

## Acknowledgements

We are thankful to: Z. Keresztes, V. Muntean, M. Alexe, A. Cristea, and A. Baricz for their technical support during sampling and sample preparation. The contribution of E. A. Levei and M. Senila in performing chemical analysis is kindly acknowledged. P-A.B was supported by the research grant PN-III-P4-ID-PCE-2016-0303 (Romanian National Authority for Scientific Research). H.B.L. was supported by the research grants: PN-III-P4-ID-PCE-2016-0303 (Romanian National Authority for Scientific Research) and STAR-UBB Advanced Fellowship-Intern (Babeş-Bolyai University). A-Ş.A. was supported by the research grants: 17-04828S (Grant Agency of the Czech Republic) and MSM200961801 (Academy of Sciences of the Czech Republic). M.M. was supported by the Postdoctoral program PPPLZ (Academy of Sciences of the Czech Republic). R.G. was supported by the research grant 17-04828S (Grant Agency of the Czech Republic).

## Contributions

H.L.B. and P-A.B. designed the study. P-A.B., A-Ş.A. and R.G. wrote the manuscript. P-A.B., A-Ş.A., R.G., M.M.S and M.M. analyzed and interpreted the data. R.G., O.B., K.I. and H.K. performed rhodopsin data analyses. M.M.S. performed CARD-FISH imaging. All authors commented on and approved the manuscript.

## Supplementary Information

### Supplementary Methods and Discussion

#### Sampling sites

Amara and Tekirghiol are naturally-formed shallow lakes, located in the South-Eastern Romania, that harbour large deposits of organic-rich sediments (or ‘sapropels’). Amara Lake (44°36.30650 N, 27°19.52950 E; 32 m a.s.l.; 1.3 km^1^ area; maximum and average depths of 6 m and 2 m respectively) is an oxbow lake with brackish water, originating from an early meander of the Ialomita river (Romanian Plain) supposedly at the end of the Neolithic Black Sea transgression (ca. 3000 BC)^2^. Tekirghiol Lake (44°03.19017 N, 28°36.19083 E; 0.8 m a.s.l.; 11.6 km^2^ area; maximum and average depths of 9 m and 3 m respectively) is a saline coastal lake derived from a marine lagoon which was isolated from the Black Sea by a narrow (˜200 m wide) sand barrier, most probably during Phanagorian Black Sea regression (ca. 500-700 BC)^1^.

#### Sediment chemical analyses

The leachable major ions were water-extracted using a sediment-to-(milli-Q) water ratio of 1:10 at room temperature. The suspension was centrifuged and the supernatant was filtered through 0.22 µm-pore sized membranes. The obtained filtrate was further analyzed for ion content (**Supplementary Table S9**). Cations (Na^+^, K^+^, Mg^2+^) were measured by inductively coupled plasma atomic emission spectrometry (ICP-AES) using Optima 5300DV spectrometer (Perkin Elmer, USA). Chloride (Cl^−^) ions were measured by titrimetric method. Sulfate (SO^2−^) was assessed by ion chromatography on ICS-1500 (Dionex, Sunnyvale, CA, USA). The analysis of salt contents of Tekirghiol and Amara sediments indicated that dominant cations and anions were (g ·Kg^−1^): Na^+^ (16.5 and 7.0), K* (1.0 and 0.22), Mg^2+^ (1.1 and 4.0), Cl^−^ (27.7 and 11.2) and SO_4_ ^2−^ (0.25 and 13.2). All chemical analyses were performed by E.A. Levei and M. Senilă at INCDO-INOE 2000 - Research Institute for Analytical Instrumentation (Cluj-Napoca, Romania).

#### Phylogenomics

The phylogenomic trees (Figure 1a, b) showed that the basal branches of Thorarchaeia were represented by MAGs recovered from the Tekirghiol hypersaline sediment (i.e. TEKIR_14 and TEKIR_12S). The other ones were shown to form a compact cluster (n = 8), which appeared to be the outcome of a more recent diversification event (as assessed by short branch lengths) in brackish environments. Thus, the two brackish Amara MAGs (AMARA_2 and AMARA_8) clustered together with the: estuarine Mai Po ones (MP8T_1, MP9T_1 and MP11T_1)^3^, Baltic Sea AB_25^4^ and the White Oak River Estuary SMTZ1_45^4^; forming a genus-level clade (as assessed by amino acid identity values). Noteworthy, this reduced phylogenomic diversity within Thorarchaeia contrasts with the highly divergent MAGs of Loki- and Heimdallarchaeia, with which it shares common ancestry (Figure 1a, b). The presence of the younger RuBisCO type (i.e. IV)^5^ (Supplementary Figure S5) in Thorarchaeia supports the ‘(more)recent diversification’, as the other Asgardaeota (sampled to date) still maintain the ancestral type III (Supplementary Figure S5).

#### Rhodopsins

The recently discovered heliorhodopsins (HeR) were reported to be present in Heimdallarchaeia^6^. HeR share low sequence similarities with all known type-1 rhodopsins, and even more remarkably they are oriented in an opposite topology in the membrane, harbouring a cytoplasmic N-terminus (in contrast type-1 rhodopsins possess extracellular N-terminus). Heimdallarchaeia appear to possess the most diverse collection of rhodopsins within the Asgardaeota superphylum at large: type-1, HeR and schizorhodopsins. In contrast Loki- and Thorarchaeia were found to contain only schizorhodopsins (SzR). Lokiarchaeia were found to harbour sequences that have a helix-turn-helix motif and which are similar to bacterioopsin-activator like proteins. These proteins, which were found to be in the proximity of the SzR (Supplementary Figure S6) have been previously characterised in *Halobacterium halobium*, where they are hypothesized to work as low oxygen sensors capable of activating the bacteriorhodopsin gene^7^. However, given the absence of the sensing N-terminal domain (NifL-like), the connection between the bacterioopsin-activator like proteins and SzR is unclear. Although, the presence of Asgardaeota in habitats with light exposure possibility (e.g. lake water column, estuarine sediments, mangrove sediments, microbial mats, etc.)^3,4^ fulfils the condition needed for rhodopsin usage, further experimental data is needed to clarify the functions of these newly described proteins.

#### Metabolism

We observed that while all Asgardaeota clades (i.e. Thor-, Loki- and Heimdallarchaeia) possess transporters for the uptake of organic compounds, their preferences towards their categories show phyla specificity. We found that the genomic repertoire of Lokiarchaeia is highly enriched (up to ten times in comparison to rest of Asgardaeota) in genes encoding for the uptake of modified monosaccharides. Thus, Lokiarchaeia had 5.85 transporters/Mb for the glycoside/pentoside/hexuronide symporters, while Thor- and Heimdallarchaeia barely reached 0.66 and respectively 0.52 transporters/Mb. We reason that this high genomic density is linked to the Lokiarchaeia’s inferred capacity to degrade cellulose (by employing the synergic action of: endoglucanases, beta-glucosidases and cellobiose phosphorylases), and to import the resulted monosaccharides into the cytosol. The newly available monosaccharides could be used to fuel the cell machinery (by transformation of glucose to acetyl-CoA or lactate with subsequent ATP production), or transformed into glycogen storage for later usage (through glycogen/starch synthases, 1,4-alpha-glucan branching enzymes; starch/glycogen phosphorylases, alpha-amylases, neopullulanases). Peptide/nickel transporters were found to have a higher density in Thorarchaeia (Thor-: 2.48 transporters/Mb; Heimdall-: 0.79 transporters/Mb; Lokiarchaeia: 0.51 transporters/Mb). As previously reported in Thor-^3,6^, we found that both Loki- and Heimdallarchaeia have the mechanisms involved in peptide and amino-acid uptake, as well as the enzymatic repertoire needed for their degradation to keto-acids and acetyl-CoA (endopeptidases: PepB, PepD, PepP and PepT; aminotransferases: AspB, ArgD, IlvE, GlmS, HisC and PuuE; glutamate dehydrogenases oxidoreductases). Among Asgardaeota, Thorarchaeia was the only phylum in which we found the glucarate and dicarboxylate uptake systems, as well as the metabolic pathways needed to catabolize putrescine to succinate. Inorganic phosphate uptake could be achieved by all Asgardaeota through the usage of PiT family transporters. Lokiarchaeia was found to harbor the PhoR-PhoB two component system involved in phosphate uptake regulation, while Heimdallarchaeia (i.e. LC_2 and RS678) was found to encode ABC-type transporters for phosphonates: refractory forms of phosphorus found to be highly abundant in marine systems^7^.

Sulfur uptake can be accounted for by the presence of sulphate permease SulP in Loki- and Thorarchaeia, as well as the ABC-type sulfonate/nitrate/taurine transport system, predominantly in the latter. Cysteine may also serve as a source of sulfur which can be mobilized by cysteine desulfurases in all three Asgardaeota phyla.

Remarkably, two Heimdallarchaeia MAGs (LC_2 and LC_3) encoded genes for the synthesis and degradation of cyanophycin (Figure 3), a non-ribosomally produced polypeptide used as a carbon and nitrogen storage pool in bacteria^8^.

While a canonical pentose phosphate pathway (PPP) is lacking in all analyzed Asgardaeota, evidence indicates that simple sugar interconversions are carried out by the non-oxidative branch of this pathway. In Thorarchaeia, the xylulose part of the non-oxidative branch was found to be largely complete with the key enzyme transketolase present in multiple MAGs. We also identified two copies of this enzyme in Heimdall_RS678, which points towards similar functional capabilities; however, with low support within the Heimdallarcheia phylum itself. The absence of transketolase enzyme, in Loki- as well as in most Heimdallarchaeia, and the presence of components of the Ribulose Monophosphate Pathway (RuMP) indicates it as a potential alternative for PPP, as previously reported in the case of *Thermococcus kodakaraensis*^9^. The presence of uridine phosphorylase within all three analyzed phyla indicates that the nucleotide degradation pathways could serve as an additional ribose source. While all analyzed Asgardaeota phyla encode components of the glycolytic pathway (i.e. type Embden-Meyerhof-Parnas), three Heimdallarchaeia MAGs (LC_3, AB_125 and AMARA_4) were found to employ non-canonical ADP-dependent kinases and, as previously noted^3^, no glucokinase homologue could be identified in Thorarchaeia. We reason that the well represented non-oxidative PPP in this group could either represent an alternative point of entry for sugars in the EMP, or that the function of canonical glucokinase is achieved by yet unidentified archaea-specific sugar kinases^10^.

Regarding pyruvate metabolism, in both Loki- and Thorarchaeia phosphoenolpyruvate (PEP) can be converted to pyruvate by phosphoenolpyruvate synthase (pps) as well as pyruvate kinase (pyk), the latter of which we identified in Heimdallarchaeia as well. Also present in all three phyla, malic enzyme (maeA) is probably responsible for catalyzing the oxidative decarboxylation of malate to pyruvate, CO_2_ and NADH. Additionally, all groups encode pyruvate phosphate dikinase (PPDK), a PPi-utilizing enzyme which interconverts PEP and pyruvate. Among the anaplerotic reactions for CO_2_ fixation, reversible carboxylation of acetyl-CoA to pyruvate may be achieved in all groups by the activity of pyruvate:ferredoxin oxidoreductase (PFOR). PEP can be synthesized by the phosphoenolpyruvate carboxykinases (PEPCK) present in all three groups, starting from oxaloacetate. Therefore, under gluconeogenic conditions, maeA and/or PEPCK, in combination with pps/PPDK, is used for directing C4 carbon intermediates from the TCA cycle, when present, to PEP^11^ - the precursor for gluconeogenesis. Noteworthy, as a hint of potential aerobiosis, we identified exclusively in Heimdallarchaeia MAGs (LC2 and LC3) genes encoding for pyruvate oxidase (poxL), which catalyzes the decarboxylation of pyruvate in the presence of phosphate and oxygen, yielding carbon dioxide, hydrogen peroxide and acetyl phosphate^12^.

Among all analyzed phyla, the complete TCA cycle was identified in Loki- and Heimdallarchaeia. Inquiringly, genomes from all three phyla, with the exception of Heimdall LC_3, were found to lack the membrane anchoring subunits of succinate dehydrogenase (sdhC, sdhD), while Lokiarchaeia contain key genes (isocitrate dehydrogenase, 2-oxoglutarate-ferredoxin oxidoreductase, ATP-citrate lyase) that are indicative of a reductive TCA cycle, involved in the autotrophic fixation of CO_2_. Also, malate dehydrogenase (MDH) was identified in Loki- and Thorarchaeia only by homology search (COG2055) and 3-dimensional structure predictions^13^.

Through analyzing new Asgardaeota genomes, we confirm previous findings^3,14^ regarding the presence of a complete Wood-Ljungdahl pathway in both Loki- and Thorarchaeia. This pathway was found absent in the partial TEKIR_3 and AMARA_4 (13 and respectively 30% CheckM^15^ estimated completeness) MAGs, as well as in the previously published (AB_125, LC_2, LC_3, RS678) Heimdallarchaeia ones. Importantly, the Wood-Ljungdahl pathway is confined to anoxic niches harboring low reducing substrates such as carbon monoxide (CO) and H ^16,17^. In spite of the fact that in anaerobic as well as aerobic microbes, CO may be used as both energy and carbon source^18^, the ability to utilize CO is conditioned by the presence of enzyme complexes known as carbon monoxide dehydrogenases (CODHs)^19^. In Heimdallarchaeia, we identified all three major subunits of the aerobic type CODH (coxSML). In this case, electrons generated from CO oxidation may be shuttled to oxygen or nitrate, which may serve as final electron acceptors^20–22^. However we observe that Thor- and Lokiarchaeia encode all components of the bifunctional and oxygen-sensitive^23^ enzyme complex known as carbon monoxide dehydrogenase/acetyl-CoA synthase (CODH/ACS). This complex is part of the Wood-Ljunghdahl pathway and is responsible for catalyzing reactions involved in autotrophic fixation of CO_2_.

Regarding oxidative phosphorylation, while V/A-type ATPase appears mostly complete in Loki- and Thorarchaeia, the other components involved in oxidative phosphorylation (the electron transport chain), are missing or incomplete, emphasizing anaerobic lifestyles. For Heimdallarchaeia we could identify complete V/A-type ATPase, succinate dehydrogenase, almost complete NADH:quinone oxidoreductase and importantly – cytochrome c oxidase – another hallmark of aerobiosis.

Uniquely among known Archaea, we identify all components of the aerobic tryptophan degradation pathway (i.e. kynurenine pathway^24^) in three published Heimdallarchaeia genomes (LC_2, LC_3, RS678). The performed evolutionary history inferences indicated that the kynurenine pathway was probably acquired by Heimdallarchaeia through lateral gene transfer from bacteria (Supplementary Figure S3). The phylogenetic trees, constructed with key enzymes of the pathway, pointed towards enzyme-specific evolutionary rates (Supplementary Figure S3). Thus, while Heimdallarchaeia MAGs formed monophyletic clusters in the dioxygenases trees (TDO, HAAO), in the kynurenine monooxygenase (KMO) one they segregated into two independent clusters. The MAG Heimdall_LC_3 showed an affinity to cluster with a sediment-dwelling Bacteroidetes (with low statistical support).

In Heimdallarchaeia the following archaellum^24^ components were identified: flaG, flaI, flaJ, flaK, flaH and a homolog of the archaeal flagellin (IPR013373). The presence of sensory methyl-accepting chemotaxis proteins (MCPs), together with a complete chemotaxis signal transduction pathway (CheA, B, C, D, R, W, Y), suggests that Heimdallarchaeia may be motile.

---

**Supplementary Figure S1.**
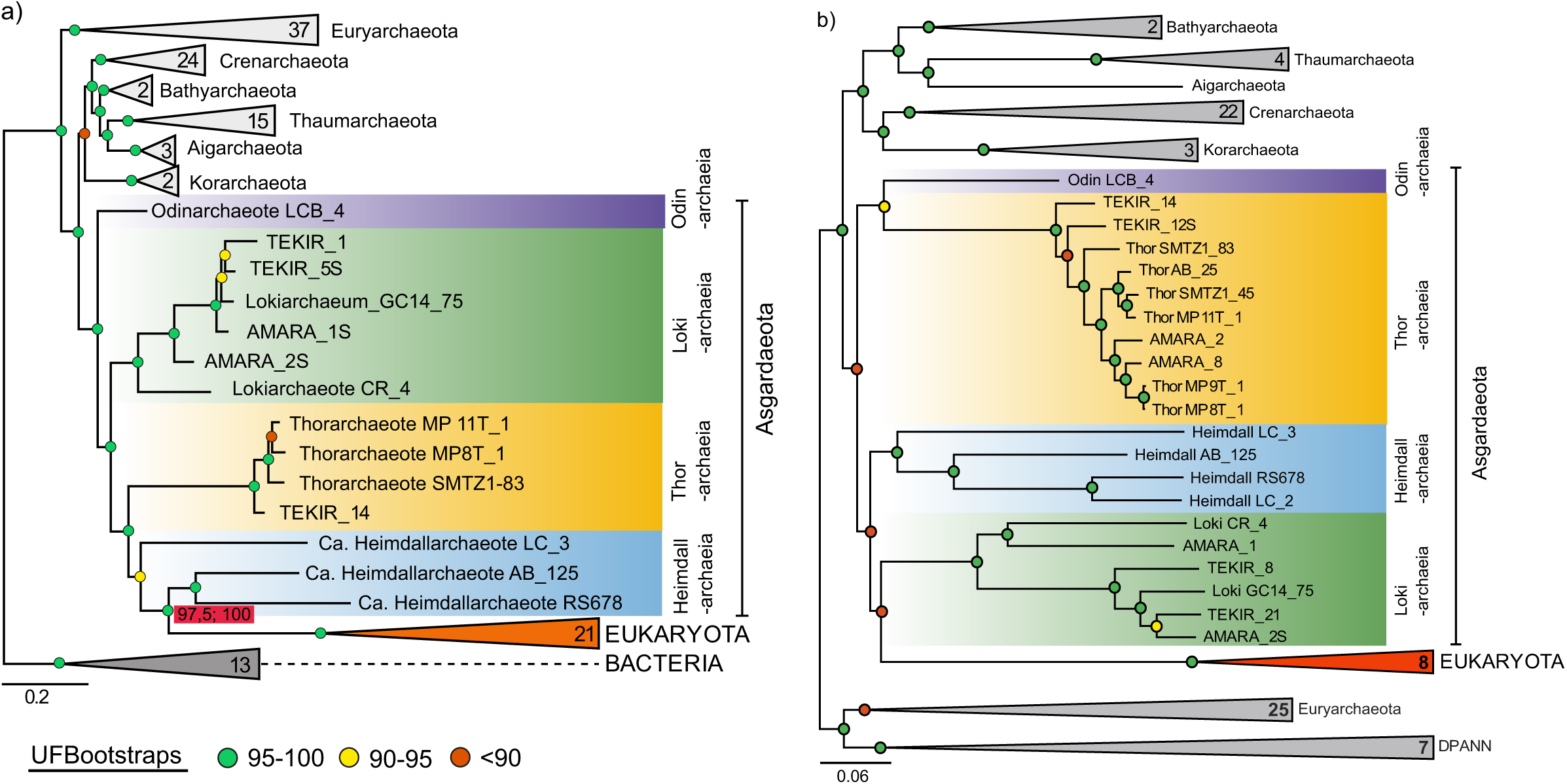
a) Maximum-likelihood phylogeny of concatenated SSU/LSU gene sequences spanning the three domains of life. The red rectangle indicates the results of the Shimodaira-Hasegawa test and ultrafast bootstrapping for the Eukaryota/Heimdallarchaeia cluster. b) Maximum-likelihood phylogenomic tree based on 48 ribosomal proteins. Bootstrap values are indicated by colored circles (legend in the lower-left part of the figure). The scale bars indicate number of substitutions per site.

**Supplementary Figure S2.**
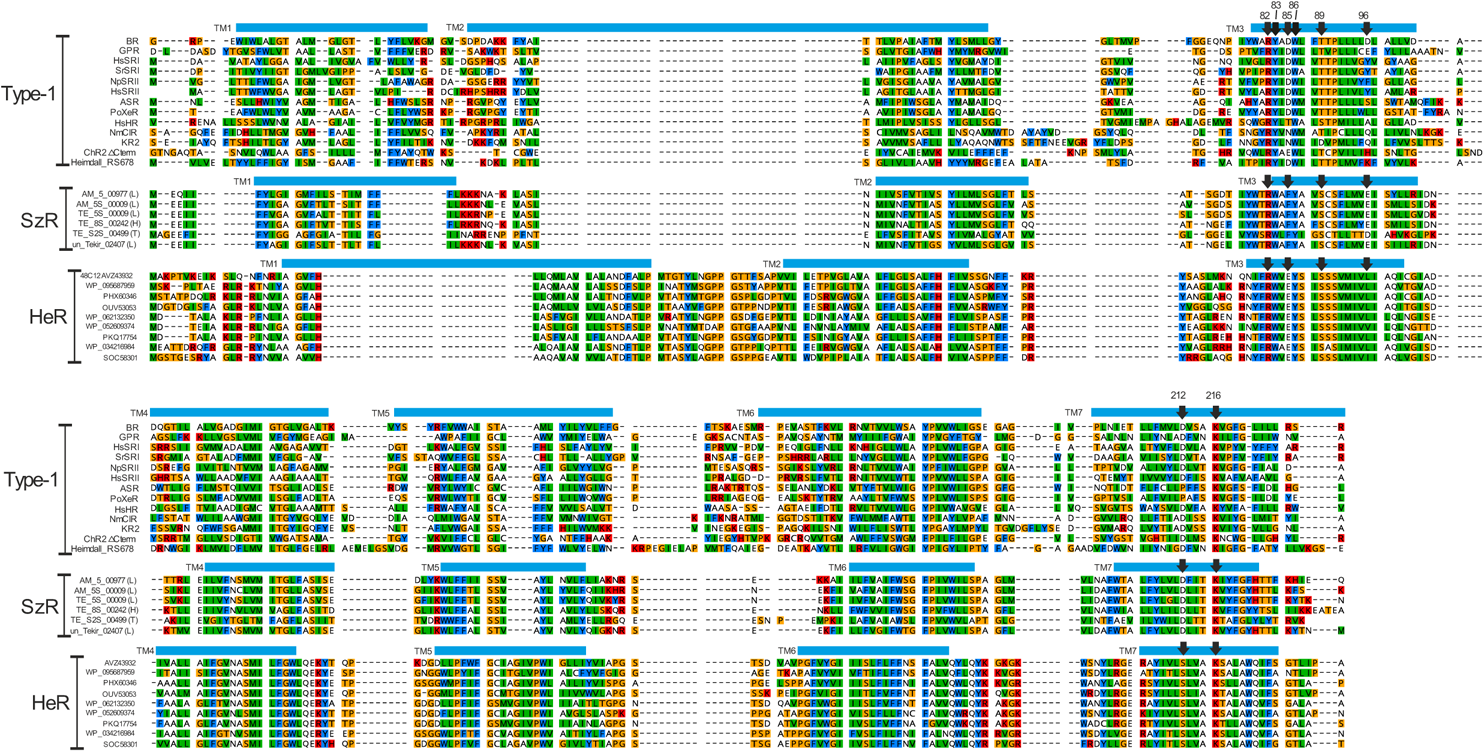
Multiple alignment of type-1, schizorhodopsins (SzR) and heliorhodopsins (HeR). Transmembrane helices (labeled as TM1-7) are shown for the first sequence of each group at the top. Sequences of type-1 rhodopsins are - BR: bacteriorhodopsin (BR), green absorbing proteorhodopsin (GPR), sensory rhodopsin I from *Halobacterium salinarum* (HsSRI) and *Salinibacter ruber* (SrSRI), sensory rhodopsin II from *Natronomonas pharaonis* (NpSRII) and *H. salinarum* (HsSRII), *Anabaena* sensory rhodopsin (ASR), xenorhodopsin from *Parvularcula oceani* (PoXeR), halorhodopsin from *H. salinarum* (HsHR), chloride-pump rhodopsin from *Nonlabens marinus* (NmClR), sodium-pump rhodopsin from *Krokinobacter eikastus* (KR2), and cation channelrhodopsin2 from *Chlamydomonas reinhardtii* (C-terminal side omitted, ChR2 ΔC-term) and putative proton-pump from Heimdallarchaeota MAG RS678 assembled from Red Sea metagenome. Sequences of Asgardaeota schizorhodopsins: Six sequences are shown, the number in brackets indicates taxonomic affiliation of the metagenome assembled genome-L – Lokiarchaeia, H – Heimdallarchaeia and T – Thorarchaeia. Sequences of heliorhodopsins are – 48C12: original actinobacterial fosmid clone from Lake Kinneret, WP_095687959: freshwater actinobacterium Ca. *Nanopelagicus abundans*, PHX60346: Actinobacteria bacterium (Lake Baikal metagenome), OUV53053: Actinomycetales bacterium TMED115 (marine metagenome), WP_062132350 Demequina aestuarii, WP_052609374: freshwater Actinobacteria bacterium IMCC26256, PKQ17754: Actinobacteria bacterium HGW-Actinobacteria-8 (groundwater metagenome) WP_034216984: *Actinoplanes subtropicus*, SOC58301: *Ornithinimicrobium pekingense*. In all groups the functionally important positions 82, 83, 85,86, 89,96, 212 and 216 (bacteriorhodopsin numbering) are marked with black arrows.

**Supplementary Figure S3.**
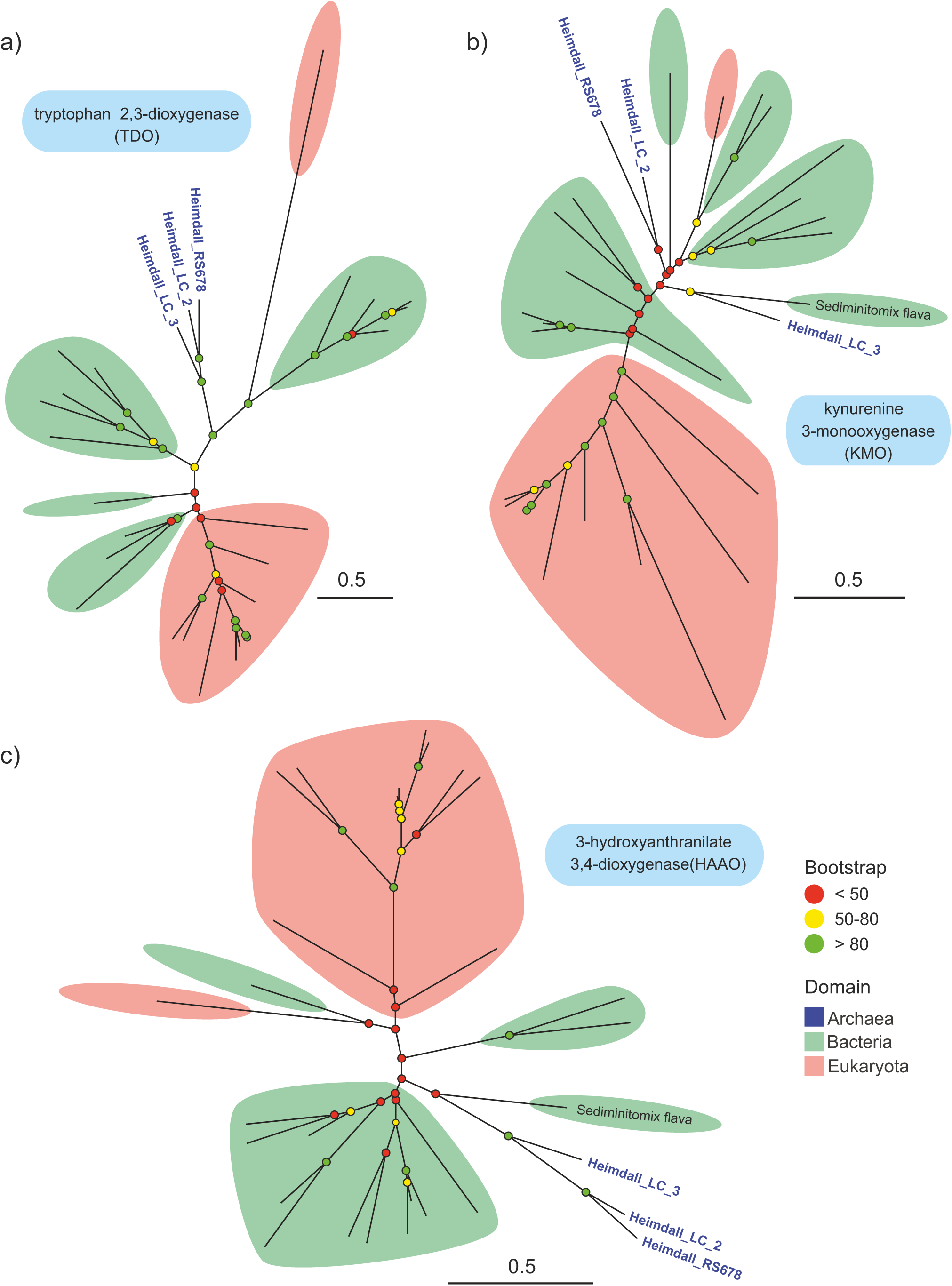
Maximum-likelihood phylogenetic trees of key enzymes of the kynurenine pathway: a) tryptophan 2,3-dioxygenase, b) kynurenine 3-monooxygenase and 3-hydroxyanthranilate 3,4-dioxygenase, respectively. Bootstrap values are indicated by colored circles (legend in the lower-right part of the figure). The names of Heimdallarchaeia MAGs are highlighted with blue color.

**Supplementary Figure S4.**
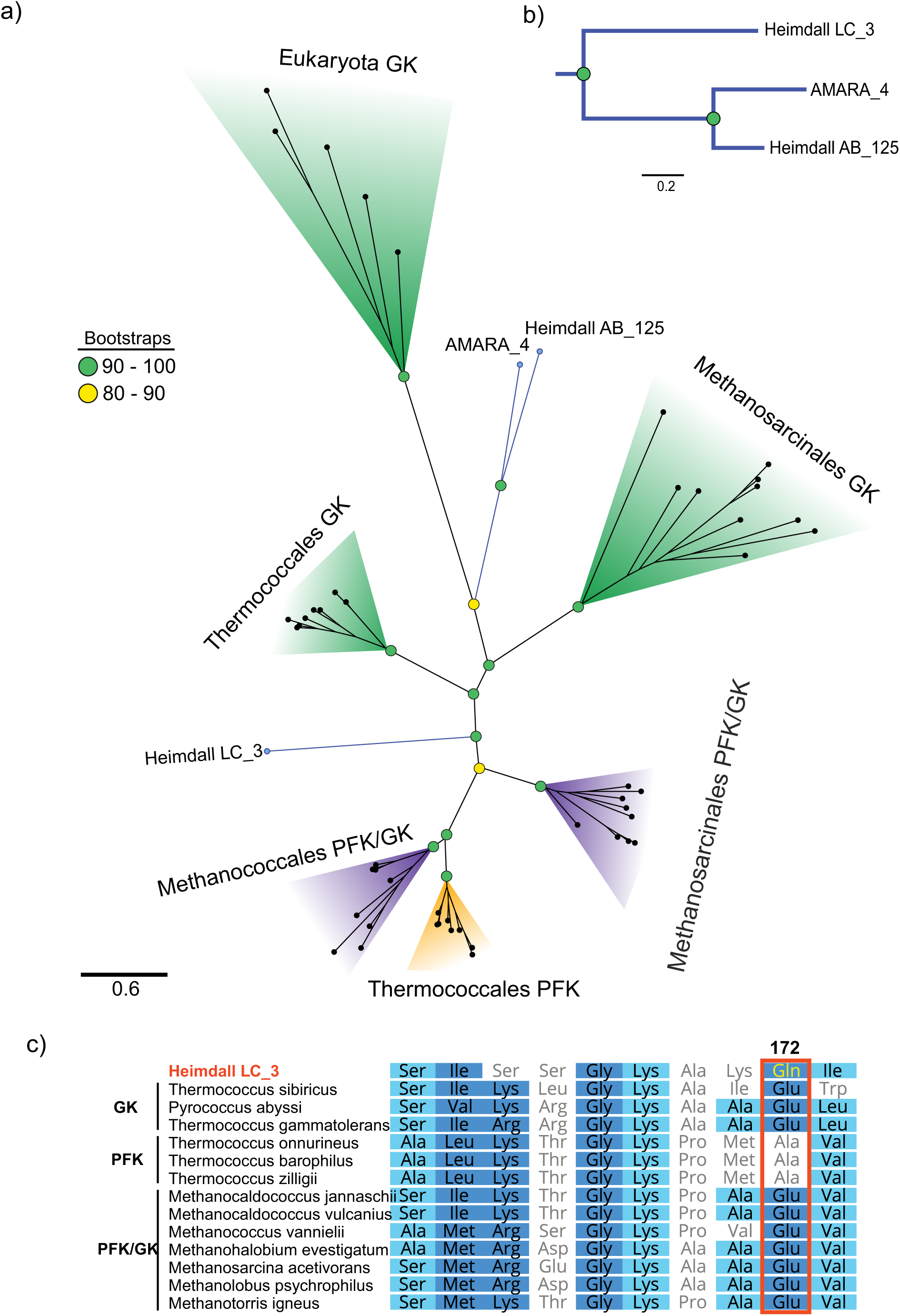
a) Maximum-likelihood phylogenetic tree of the the ADP-dependent kinases family in Archaea and Eukaryota. The branches belonging to Heimdallarchaeia are colored in blue. The colored panels highlight clades with: glucokinase activity-green, phosphofructokinase activity -orange, and both gluco- and phosphofructokinase activity-purple. b) Phylogenomic subtree, generated using maximum-likelihood methods, which shows Heimdall_LC_3 MAG as the oldest within the ones that were found to harbor ADP-dependent kinases. The scale bars indicate number of substitutions per site. Bootstrap values are indicated by colored circles (legend in the lower-right part of the figure). c) Sequence alignment of ADP-dependent kinases highlighting the functionally important position 172, as inferred from Castro-Fernandez et al. 2017. The similarity-coloring scheme is based on the BLOSUM62 matrix.

**Supplementary Figure S5.**
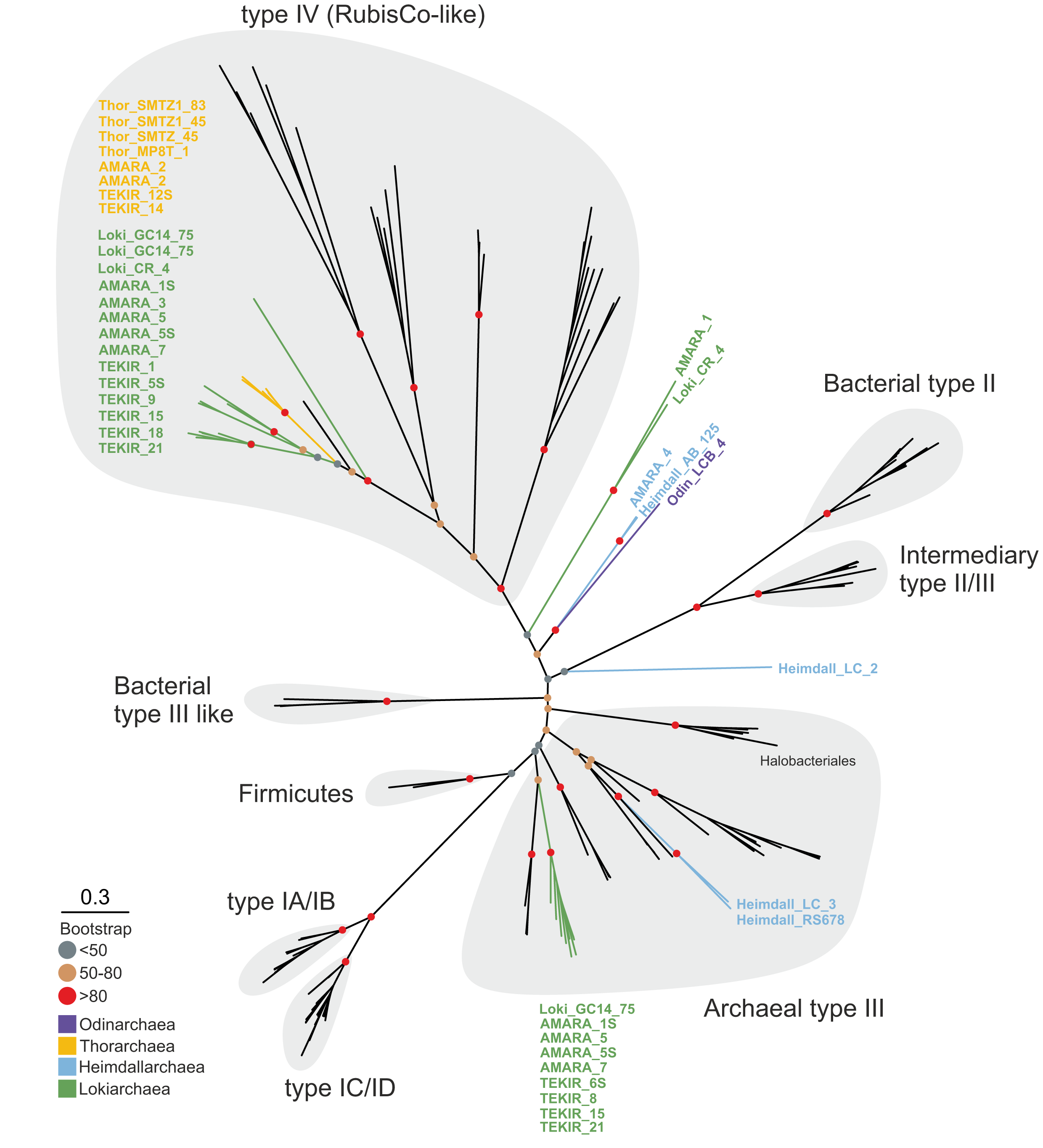
Maximum likelihood tree of the large subunit of RubisCo (types I-III) and RubisCo-like (type IV) (rbcL, K01601) protein sequences (n=146) of bacterial and archaeal taxa. Reference sequences were chosen based on previous trees constructed by Tabita et al. 2007 and Wrighton et al. 2016. The branches representing the members of Asgard superphylum are colored based on their affiliated phylum (legends on the bottom left). The scale bar indicates the amino acid substitutions per site.

**Supplementary Figure S6.**
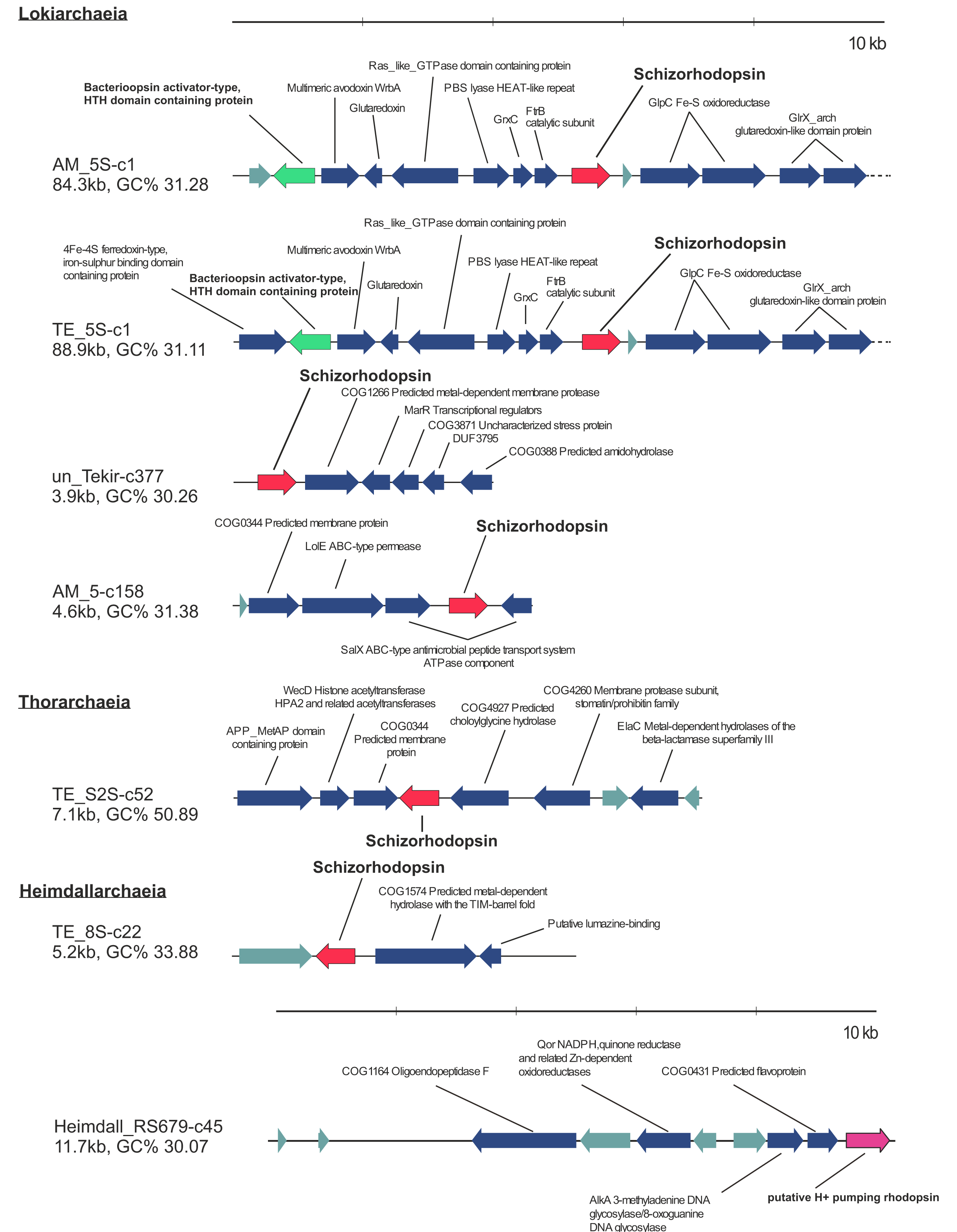
Genomic context of rhodopsins in Asgardaeota MAGs. Schizorhodopsins are shown in red, bacteriorhodopsin activator-like protein in green, bacteriorhodopsin in purple and the hypothetical proteins in grey. 10 kb scale bars are shown at the top of each category.

**Supplementary_Table_S1.xlsx**

This file contains general statistics for MAGs recovered from the Amara and Tekirghiol Lakes. Estimated genome length (EGS) was determined for MAGs with completeness >70%.

**Supplementary Table S2.**
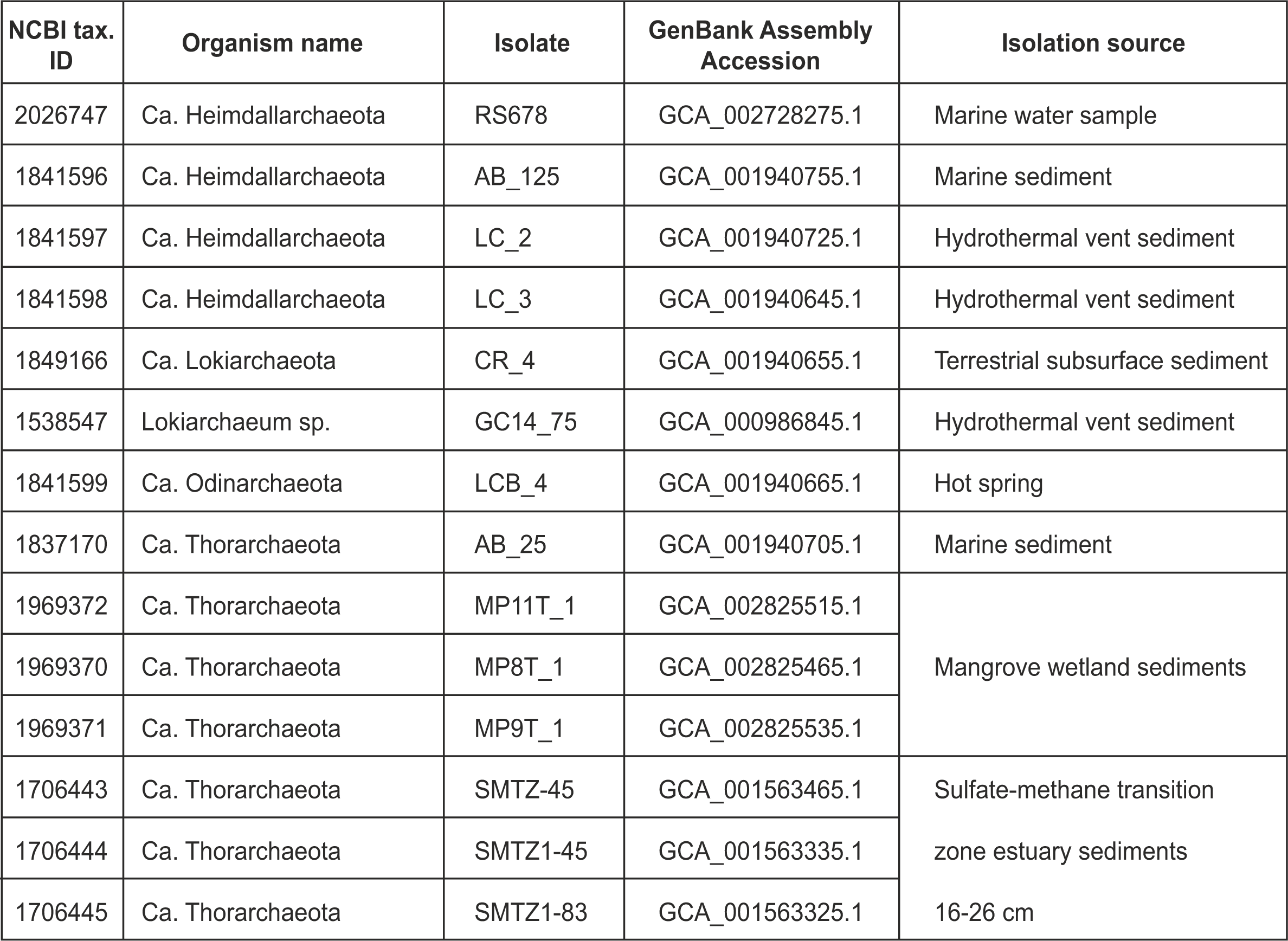
List of Asgardaeota genome assemblies downloaded from the NCBI genomes repository (https://www.ncbi.nlm.nih.gov/genome) for this study.

**Supplementary_Table_S3.xlsx**

KEGG orthology annotation for Asgardaeota MAGs.

**Supplementary Table S4.** Accession numbers list for INTERPRO (IPR) domains, COGs (Cluster of Orthologous Groups) and UniProtKB protein sequences used to identify potential Eukaryotic Signature Proteins in Asgardaeota MAGs. Particularly, the first ESP (1*) was identified by local scanning in MAGs with BLASTP using human sequences downloaded from UNIPROT (https://www.uniprot.org/).

| Nr. crt. | Description | IPR domain/COG/UniProtKB |
| --- | --- | --- |
| 1* | DNA polymerase, ε-like catalytic subunit | Q59EA9, Q9UNF3, Q9Y5S5, Q07864, Q9Y5S4, F5H1D6 |
| 2 | Topoisomerase IB | arCOG08649, COG3569 |
| 3 | RNA polymerase, subunit G (rpb8) | IPR031555 |
| 4 | Ribosomal protein L22e | IPR002671 |
| 5 | Ribosomal protein L28e/Mak16 | IPR029004 |
| 6 | Tubulins | IPR000217 |
| 7 | Actin family (divergent) | IPR004000 |
| 8 | Actin/actin-like conserved site | IPR020902 |
| 9 | Arp2/3 complex subunit 2/4 | IPR008384 |
| 10 | Gelsolin-domain protein | IPR007122 |
| 11 | Profilin | IPR036140 |
| 12 | ESCRT-I: Vps28-like | IPR007143 |
| 13 | ESCRT-I: steadiness box domain | IPR017916 |
| 14 | ESCRT-II: EAP30 domain | IPR007286 |
| 15 | ESCRT-II: Vps25-like | IPR014041 and IPR008570 |
| 16 | ESCRTIII: Vps2/24/46-like | IPR005024 |
| 17 | Ubiquitin-domain protein | IPR000626 |
| 18 | E2-like ubiquitin conjugating protein | IPR000688 |
| 19 | E3 UFM1-protein ligase 1 | IPR018611 |
| 20 | RAG-type GTPase domain | IPR006762 |
| 21 | Longin-domain protein | IPR011012 |
| 22 | Vacuolar fusion protein Mon1 | IPR004353 |
| 23 | RLC7 roadblock domain protein | IPR004942 |
| 24 | TRAPP-domain protein | IPR007194 |
| 25 | Zinc finger, Sec23/Sec24-type | IPR006896 |
| 26 | Arrestin-like proteins (C-terminal) | IPR011021 |
| 27 | Vesicle coat complex COPII sub. SEC24/SFB2/SFB3 | COG5028 |
| 28 | Folliculin (N-terminal) | IPR037520 |
| 29 | Ribophorin I | IPR007676 |
| 30 | Oligosaccharyl transf. OST3/OST6 | IPR021149 |
| 31 | Oligosaccharyl transf. STT32 | IPR003674 |
| 32 | Ezrin/radixin/moesin C-terminal domain | IPR011259 |
| 33 | active zone protein ELKS | IPR019323 |

**Supplementary_Table_S5.xlsx**

List of taxa used for phylogenetic ribosomal RNA (small subunit SSU and large subunit LSU) as well as ribosomal protein-based inferrences based mainly on a previously published list https://www.nature.com/articles/nature21031. Silva SSU and LSU rRNA IDs are indicated. For MAGs recovered in this study, contig ID and positions are indicated (highlighted).

**Supplementary Table S6.**
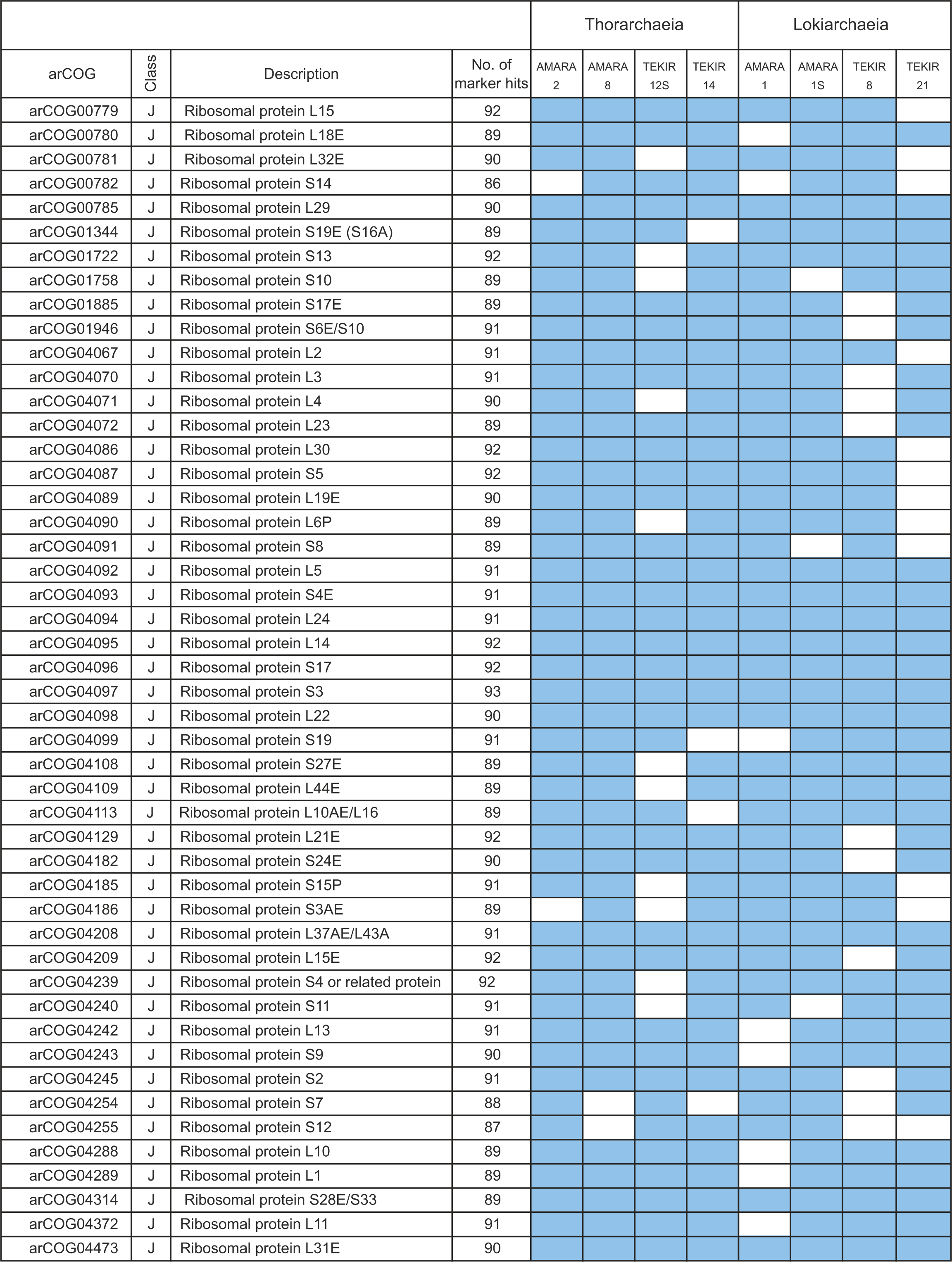
List of the 48 ribosomal proteins used for phylogenomic analyses. Total number of taxa (from the 93 included) in which a particular marker was identified is indicated as ‘No. of marker hits’. Blue/white cells indicate presence/absence respectively.

**Supplementary Table S7.**
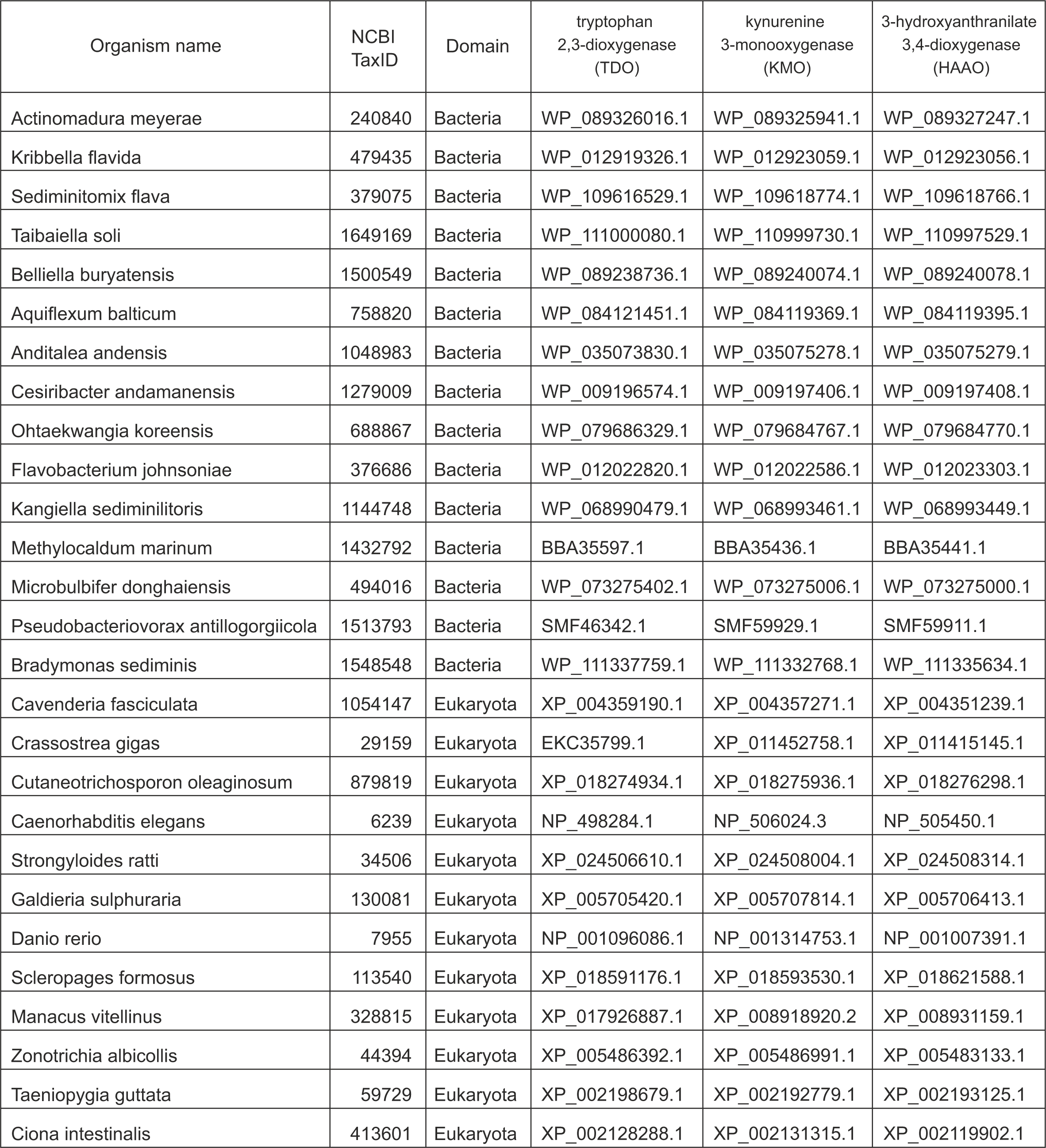
Protein RefSeq accession numbers for sequences used in phylogenetic inferences for key components of the Tryptophan degradation pathway (kynurenine pathway) in Heimdallarchaeia.

**Supplementary Table S8.**
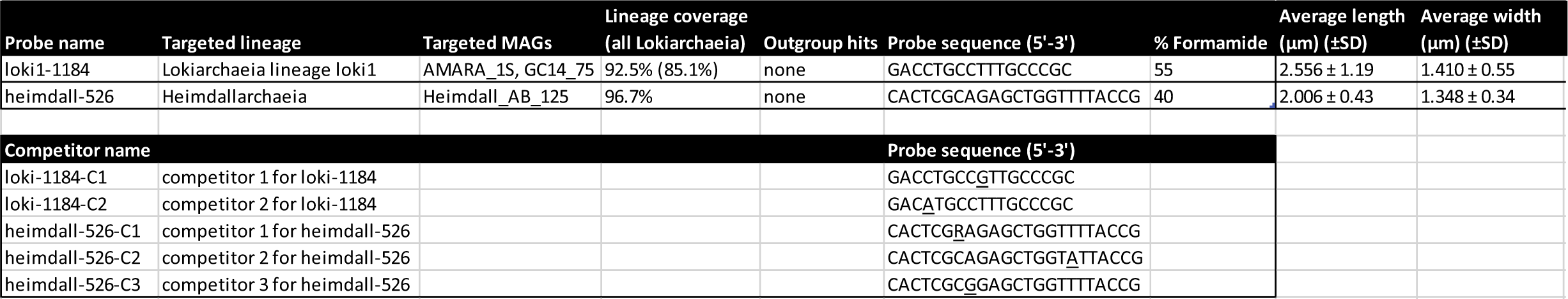
Probes designed for CARD-FISH.

**Supplementary Table S9.** Major water leachable ions and elements from sediments collected in Tekirghiol and Amara Lakes during October 2017.

| **Ions/metals**<br>**(mg·Kg<sup>-1</sup>)** | **Tekirghiol**<br>**Lake** | **Amara**<br>**Lake** |
| --- | --- | --- |
| pH of pore water | 7.38 | 8.03 |
| Na<sup>+</sup> | 16520 | 7000 |
| K<sup>+</sup> | 1020 | 221 |
| Ca<sup>2+</sup> | 86.4 | 640 |
| Mg<sup>2+</sup> | 1100 | 4000 |
| Fe<sup>*</sup> | 0.45 | 0.38 |
| Mn<sup>*</sup> | <0.1 | 0.57 |
| Cl<sup>-</sup> | 27700 | 11200 |
| SO<sub>4</sub><sup>2-</sup> | 250 | 13240 |

**Note**: Values are determined for the water-extracted fraction of the wet sediment. The * indicates element total concentration.

